# Decoupling of stomatal conductance from net assimilation at high temperature as a mechanism to increase transpiration

**DOI:** 10.1101/2025.11.03.686201

**Authors:** Philipp Schuler, Margaux Didion-Gency, Giovanni Bortolami, Thibaut Julliard, Günter Hoch, Christoph Bachofen, Ansgar Kahmen, Charlotte Grossiord

## Abstract

- Photosynthetic assimilation (A_net_) and stomatal conductance (g_s_) are usually strongly coupled, but this relationship is decreased or even lost at high temperatures (T_air_). The contributions of environmental drivers (T_air_, vapour pressure deficit (VPD), and soil moisture) in interaction with the physiological mechanisms behind this process are still unclear.
- We exposed saplings of three temperate and tropical species to rising T_air_ (20 to 40°C) at low (1.2 to 1.9 kPa) and increasing VPD (1.1 to 5.6 kPa), and at stable T_air_ (35°C) to increasing VPD (1.4 to 4.3 kPa) under well-watered or chronic soil drought conditions (≤10 %). A_net,_ g_s_, and transpiration (E) in the light and the dark and leaf thermoregulation were tracked throughout the experiment.
- When VPD remained low, g_s_ continued to increase while A_net_ decreased at T_air_ > 35°C, leading to stomatal decoupling. In contrast, under rising VPD, trees maintained the coupling between A_net_ and g_s_ at high T_air._
- While a decoupling of A_net_ and g_s_ only occurred when VPD was low, A_net_ and E decoupled under both VPD regimes at high T_air_.
- Our results indicate that, since g_s_ and VPD collectively drive E, stomatal decoupling is needed to increase E when VPD is not sufficiently high.

## Introduction

It is widely assumed that net photosynthesis (A_net_) and stomatal conductance (g_s_) are optimally coupled to maximize carbon gain while minimizing water loss (Wong et al., 1979; Lawson et al., 2010; Duursma et al., 2014). Hence, when A_net_ declines at high air temperatures (T_air_) due to biochemical limitations (Sage & Kubien, 2007), g_s_ should decrease simultaneously (Wong et al., 1979; Lawson et al., 2010; Duursma et al., 2014). Most leaf-level stomatal models assume this strong coupling (e.g., Farquhar & Wong, 1984; Medlyn et al., 2011). Yet, more and more studies are challenging this understanding, both in natural and controlled conditions, showing a decoupling (i.e., the loss of the strong relationship) between A_net_ and g_s_ under high T_air_ (Schulze et al., 1973; Aphalo and Jarvis, 1991; Eamus et al., 2008; Urban et al., 2017; Schönbeck et al., 2022; Marchin et al., 2023; Diao et al., 2024a; Gauthey et al., 2024) – a process referred to as “stomatal decoupling”. In most studies, while A_net_ tends to drop at T_air_ above 30-35°C (i.e., reflecting the optimum Tair for photosynthesis), g_s_ either remains stable (e.g., Schönbeck et al., 2022; Gauthey et al., 2024) or rises (e.g., Schulze et al., 1973; Urban et al., 2017; Diao et al., 2024a). Keeping high g_s_ (and hence transpiration, E) could be a mechanism that serves to avoid lethal thermal damage during heatwaves through continued evaporative cooling (e.g., Drake et al., 2018; Gauthey et al., 2024, Bachofen et al., 2025), even though it increases the risk of embolism (e.g., Schoenbeck et al., 2022). However, how high T_air_, high vapor pressure deficit (VPD), and low soil moisture - factors that usually co-occur during heatwaves - contribute to the decoupling of A_net_ and g_s_ remains unclear. Similarly, the species-specific variability and the physiological mechanisms behind stomatal decoupling at high T_air_ remain elusive (Mills et al., 2024), leading to significant uncertainties in leaf-level stomatal models and our understanding of tree thermal tolerance.

Potential drivers of the stomatal decoupling at high T_air_ can be separated into purely physical and physiological components (see Mills et al., 2024, for a detailed review of these processes). Physical drivers include the T_air_-driven increase in the diffusive rate of water vapour in the air (Massman, 1998), which leads to increasing g_s_ at increasing T_air_ even if the opening of the stomatal aperture remains constant. This is considered the baseline for the observed stomatal decoupling. In addition, higher T_air_ will increase the xylem hydraulic conductivity (K_leaf_) because water viscosity declines sharply with increasing T_air_ (Matzner and Comstock, 2001), leading to less negative leaf water potential (Ψ) for a given transpiration rate, and hence, higher g_s_ (Mills et al., 2024). Higher T_air_ will also raise the minimum epidermal conductance (i.e., residual water loss once stomata are closed, g_min_) (Duursma et al., 2014), which could further contribute to the higher measured g_s_ with warmer air (Gauthey et al., 2024). However, a strong T_air_-driven increase of g_min_ likely only occurs at high T_air_ > 40°C (Wang et al., 2024). Moreover, while photosynthesis, to our current understanding, does not directly impact the stomatal apparatus (Von Caemmerer et al., 2004; Rogers et al., 1980; Urban et al., 2017), it might have an indirect influence via T_air_-driven shifts in photosystem (PS) I and PS II activity (Messinger et al., 2006). High T_air_ alters the balance between the proton generation in PS II and the proton consumption in PS I, causing changes in the redox state inside the cells (Busch, 2014). Increasing photorespiration at high T_air_ (Dusenge et al., 2019) or degradation of Rubisco above 40°C (Bose et al., 1999) might be reasons for an altered ratio between PS I and PS II. Such changes in the redox state could be sensed by guard cells (Lawson et al., 2010), therefore influencing the stomatal aperture and increasing g_s._ Together, these physical and physiological processes establish a foundational framework through which rising T_air_ alone could drive increases in g_s_, independent of changes in stomatal aperture or carbon assimilation. Still, most mechanisms have never been tested experimentally.

In addition, stomatal decoupling should depend on the environmental conditions experienced by the plants, particularly the evaporative demand and soil moisture (Schulze et al., 1973; Mott and Peak, 2010). An increase in VPD (e.g., Diao et al., 2024b) and a reduction in soil moisture (e.g., Brodribb and Holbrook, 2003) trigger a decline in leaf Ψ, lowering the vapour content in the stomatal pore, which in turn leads to stomatal closure (Peak and Mott, 2011), reduced A_net_ and g_s_ (Grossiord et al., 2020), and may prevent stomatal decoupling. Yet, multiple studies have reported stomatal decoupling during natural heatwaves where elevated T_air_ co-occurs with high VPD and low soil moisture (e.g., Drake et al., 2018; Marchin et al., 2023; Gauthey et al., 2024), challenging these observations and our understanding of gas exchange regulation. Moreover, while drought typically suppresses E through strong stomatal closure (Hall and Schulze, 1980; Bachofen et al., 2023), increasing VPD can enhance E due to a greater evaporative demand (Massmann et al., 2019), until stomatal limitations dominate under severe stress. Further complicating our understanding of g_s_ and E responses to high T_air_, the decoupling appears highly species-specific. Among the 22 species reviewed by Mills et al. (2024), only half exhibited consistent stomatal decoupling, with varying intensity—on average, gₛ doubled between 10°C and 40°C. Nevertheless, given the limited number of species studied, the extent to which this pattern applies across taxa remains uncertain. Whether or above which T_air_ threshold stomatal decoupling occurs probably differs between plants with different climatic adaptations (Mills et al., 2024). Species adapted to low soil moisture and relatively high VPD, such as in many temperate regions, likely respond differently than species growing in high soil moisture and low VPD environments, such as in the wet tropics (Cunningham, 2004; Middleby et al., 2024; Slot et al., 2024). However, no study has explored the variation in stomatal decoupling across species originating from contrasting climatic types, the relative influence of T_air_, VPD, and soil moisture in regulating this process.

We hypothesized that (1) stomatal regulation at high T_air_ will be strongly impacted by VPD and soil moisture availability, with reductions in g_s_ at high VPD and in dry soil, while it would increase with T_air_ at stable VPD and moist soil conditions (i.e., leading to stomatal decoupling and increased E). Further, (2) stomatal regulation of tree species from temperate regions should be more sensitive to high T_air_, thereby increasing decoupling at high T_air_, low VPD, and moist conditions. In contrast, tropical species, which evolve at higher T_air_ and VPD environments, may show a lower magnitude of decoupling under the same conditions. Therefore, (3) we expect stomatal decoupling to contribute to leaf temperature regulation, with higher E leading to stronger evaporative leaf cooling at higher T_air_ for temperate and tropical tree species.

## Materials and methods

### Plant handling prior to the experiment

In January 2024, approximately 50-100 cm tall saplings of three tree species from the European temperate climates (Köppen climate classification: *Alnus cordata* (Loisel.) Duby (Cfa-Cfb), *Acer platanoides* L. (Cfa), *Phillyrea angustifolia* L. (Csa) – 30 replicates per species) and three tree species from tropical wet to monsoonal climates (*Terminalia microcarpa* Decne. – 30 replicates, *Trema tomentosa* (Roxb.) H. Hara, *Syzygium jambos* L. (Alston) – 24 replicates each (all three species from Am-Af climates)) were planted in 5L pots, using Oekohum “Container CLASSIC – ohne Torf” (ökohum gmbh, 8585 Herrenhof, Switzerland), a mixture of bark humus, bark compost, wood fiber, hemp fiber, coconut, miscanthus, expanded clay, perlite. The species were selected to represent different phylogenetic lineages (Table S1).

Prior to the experiment, the tropical tree species were grown in a greenhouse with a T_air_ of 20-25°C and a relative humidity (RH) between 50-60%. All trees were transferred to a greenhouse with additional overhead lights (10 hours a day), where the T_air_ was kept at 25°C and the RH at 50% (stable VPD of 1.6 kPa) until they were exposed to the experimental conditions. The flushing of the deciduous temperate tree species started within 2 weeks, and for *P. angustifolia*, after about one month, and their leaves were fully mature when they were used for the experiment.

The six species were exposed sequentially to the experimental conditions, which lasted 12 days. One week before the experiment, we reduced watering to half of the randomly selected saplings for the drought treatment to ensure low soil moisture. Soil moisture was measured every second day using a handheld soil moisture probe (TDR100 Soil Moisture Meter, FieldScout, Hoskin Scientific, Burnaby, Canada).

Two days before the experimental conditions started, each tree species was transferred into the three climate chambers (see experimental details below). During these two days, the trees were acclimated to the conditions in the climate chamber at a T_air_ of 25°C, an RH of 50%, a day/night cycle of 12/12 hours, and an average photosynthetic active radiation (PAR) of ∼700 µmol m^-2^ s^-1^ (no UV radiation). During the acclimation period, the T_air_ was decreased by 5°C during the night, but VPD was held constant.

### Experimental design

To disentangle the individual and additive effects of T_air_, VPD, and soil water availability on the measured parameters (see below), the plants were exposed to three different T_air_ and VPD regimes, each with a subset of well-watered or drought conditions (n=5 trees per species, atmospheric regime, and soil moisture treatment; n=4 for *T. tomentosa* and *S. jambos*). The setpoints for the three T_air_ and VPD regimes were:

1. “increasing T_air_, low VPD”: increasing T_air_ setpoints from 20 to 40°C (20.0°C, 25.0°C, 30.0°C, 32.5°C, 35.0°C, 40.0°C) and maintaining a low VPD around 1.2 kPa by increasing setpoints RH (50%, 62%, 72%, 75%, 79%, 85%, respectively).
2. “increasing T_air_ and VPD”: rising T_air_ from 20 to 40°C as in (1) but increasing VPD by decreasing RH setpoints (50%, 37%, 28%, 24%, 21%, 16%; VPD = 1.2 kPa, 2.00 kPa, 3.1 kPa, 3.7 kPa, 4.4 kPa, 6.2 kPa, respectively).
3. “stable T_air_, increasing VPD”: constant T_air_ at 35°C while increasing VPD by reducing RH setpoints (80%, 65%, 55%, 40%, 30%, 20%).

The actual average T_air_ and VPD of the chambers can be found in Table S3. Each of the six T_air_/VPD steps was maintained for two days, with the measurements (detailed below) conducted systematically on the second day to ensure plants had been exposed to the corresponding conditions. As during the acclimation period, T_air_ was decreased by 5°C during the night (relative to the previous daytime conditions), but VPD was held constant during day- and nighttime.

The mean soil moisture (soil volumetric water content, VWC) was 32.6% ± 0.3% SE, 32.4% ± 0.4% SE, and 31.2% ± 0.4% SE (“increasing T_air_, low VPD”, “increasing Tair and VPD”, and “stable T_air_, increasing VPD”, respectively) for well-watered plants and 9.1% ± 0.3% SE, 9.2% ± 0.3% SE, and 7.4% ± 0.3% SE (“increasing T_air_, low VPD”, “increasing T_air_ and VPD”, and “stable T_air_, increasing VPD”, respectively) for drought-exposed plants (Table S2). While the soil moisture of the well-watered plants showed no significant trend during the experimental period, drought-exposed plants had a significant decrease in soil moisture from 12.2 ± 0.7% SE to 7.1% ± 0.5% SE, from 12.7 ± 0.9% to 6.6% ± 0.6%, and from 11.7% ± 0.8% SE to 5.6% ± 0.2% SE for “increasing T_air_, low VPD”, “increasing T_air_ and VPD”, and “stable T_air_, increasing VPD”, respectively (Table S2).

### Leaf-level measurements on light- and dark-adapted leaves

To determine the importance of stomatal decoupling, as well as potential underlying drivers, leaf-level gas exchange in the light (A_net_, E, g_s_) and dark (g_s dark_, E_dark_), and T_air_ (inside the Li-6800 measuring chamber) as well as leaf temperature (T_leaf_) were measured every second day (i.e., at days 2, 4, 6, 8, 10, 12) with a Li-6800-01A (LI-COR Environmental, Lincoln, NE 68504, United States). In the morning, one light-exposed leaf from the upper part of the canopy was dark-adapted by covering it in aluminium foil for more than 20 minutes. A second leaf next to the one selected for the dark-adapted measurements was chosen for the light-adapted ones. The dark-adapted leaves were measured with the following settings: the cuvette Tair and RH were set to the current values of the corresponding climate chamber, a reference CO_2_ concentration of 400 ppm, a PAR of 0 µmol m^-2^ s^-1^, and a flow of 500 µmol s^−1^. After every dark-adapted measurement, the light-adapted measurements had the same settings, except that PAR was set to 1500 µmol m^-2^ s^-1^, ensuring light saturation for all species. All measurements were taken once the gas exchange stabilized, typically after 5-10 minutes. The average T_air_ and VPD conditions during the measurements can be found in Table S3.

### Data analysis

The collected data was analyzed and displayed in R v.4.2.0 (R Core Team, 2025) and base R. WUE_i_ was calculated by dividing A_net_ by g_s_, WUE was calculated by dividing A_net_ by E, and the ratio between g_s_ and g_s dark_ (g_s_-ratio) as well as E and E_dark_ (E-ratio) was calculated by dividing the former by the latter. Furthermore, we calculated the difference between T_air_ and leaf T_leaf_ (T_offset_).

To reduce “noise” due to species-specific offsets and display the general response of temperate and tropical species to T_air_ and VPD, we normalized the A_net_, g_s_, g_s dark_, E, and E_dark_ to their species-specific average values per treatment of day 2. However, not for the analysis of the relationship between any of these, and also not to calculate WUE_i_, WUE as well as the g_s_-ratio and E-ratio). Data from day 8 (T_air_ = 32.5°C) were excluded from the analysis to maintain consistent 5°C steps.

Overall responses of A_net_, g_s_, g_s dark_, g_s_-ratio, E, E_dark_, E-ratio, WUE_i_, WUE, T_offset_, and T_offset dark_ to T_air_ and VPD were displayed in simple ggplots plots with LOESS smooths without stats by using ggplot2 to display the data in plots (H Wickham, 2016).

To statistically analyze the responses to T_air_ and VPD for the subsets (“increasing T_air_, low VPD”, “increasing T_air_ and VPD”, and “stable T_air_, increasing VPD”, split into temperate/tropical, wet/dry), we modeled the relationship between A_net_, g_s_, g_s dark_, E, E_dark_ (all normalized), WUEi, WUE. Toffset, T_offset dark_, and T_air_ with ordinary least squares (stats::lm) and then tested for a significant slope change (breakpoint) using the segmented (Fasola et al., 2018) package’s Davies test; when indicated, we fit a one-breakpoint segmented regression, and extracted R², p-values, estimated breakpoint(s), and segment slopes. Figures were produced with ggplot2, ggpmisc (Aphalo, 2025), ggpubr (Kassambara, 2025), patchwork (Pedersen, T., 2025) for layout/annotations.

Furthermore, we tested the relationship between A_net_ and g_s_, A_net_ and E, E and g_s_, whether g_s_ and E predicted T_offset_, both in the light and the dark, using a linear model with T_air_ (°C) (for the two conditions with increasing T_air_) or VPD (kPa) (for the condition with stable T_air_) as a categorical factor and, for instance, g_s_ x T_air_ interactions (lm(A_net_ ∼ g_s_ * T_air_)). The simple slopes at each T_air_ or VPD, respectively, were estimated with estimated marginal trends, pairwise-compared and compact-letter-displayed (cld) using the R package “emmeans” (Lenth and Piaskowski, 2025), and we reported slope estimates with p-values. The impact of E and g_s_ on T_offset_ was done in the light and the dark to be able to observe other potential impacts of leaf traits, such as different absorbance or other leaf properties, on the efficiency of transpirational leaf cooling. Lastly, to assess the overall relationships between g_s_ and E with T_offset_ in the light and the dark, we fitted separate simple linear regressions per T_air_ or VPD (T_offset_ ∼ E, etc.) across the individual subsets to report R² and slope significance (with significance codes *, **, ***).

## Results

### T_air_ and VPD response of net assimilation

Across experimental conditions, net assimilation (A_net_) responses to T_air_ and VPD were highly variable between biogeographic groups and soil water availability (Figs. 1; S1, S2).

**Figure 1:**
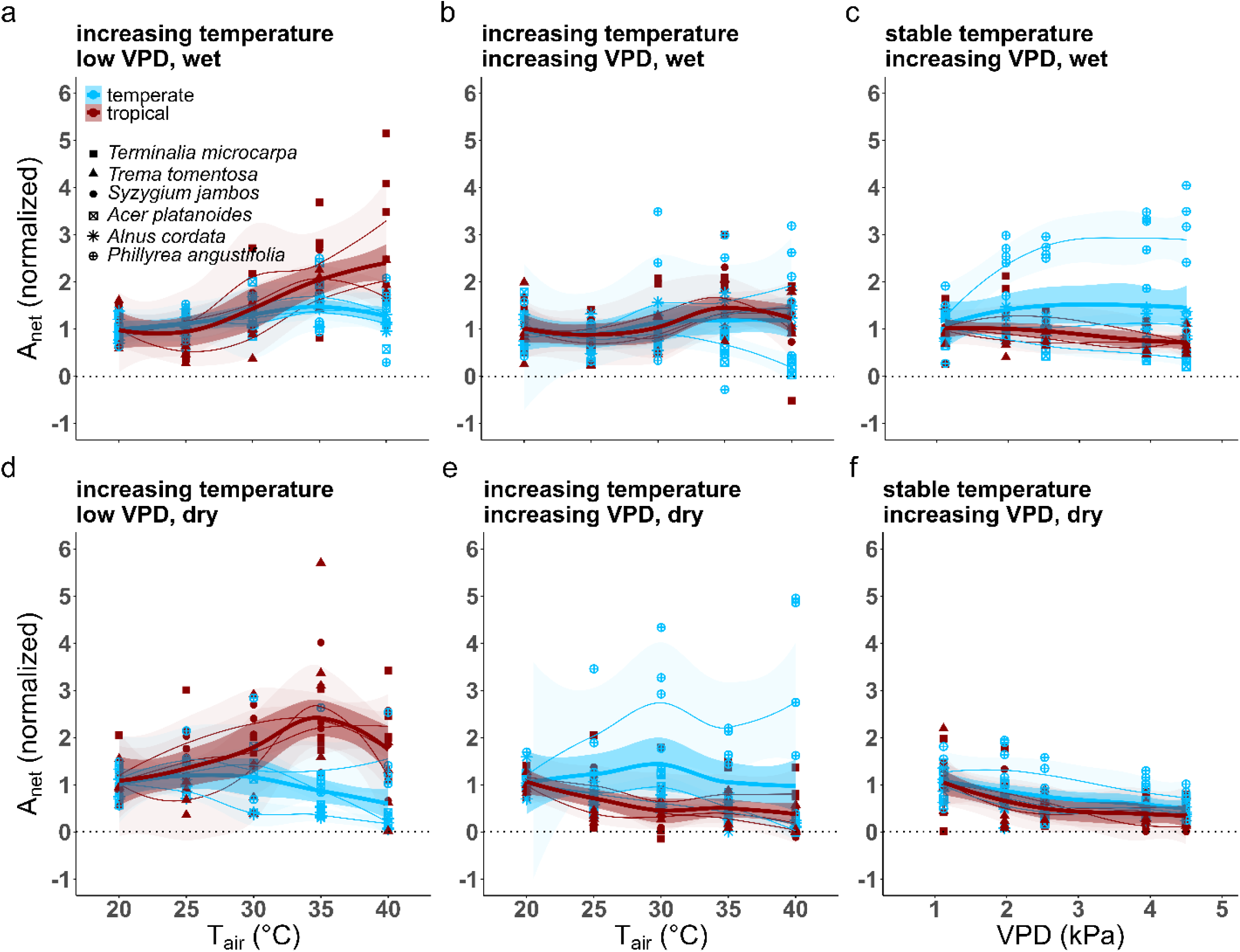
Response of the normalized net assimilation (A_net_, normalized to the species-specific average value at day 2) when temperature (T_air_) increases while vapour pressure deficit (VPD) remains low (a, d), when T_air_ and VPD simultaneously increase (b, e), and when VPD increases at a stable T_air_ of 35°C (c, f) of well-watered plants (a, b, c) or drought-exposed (d, e, f) plants. Points show individual measurements, with color indicating biogeographic origin (temperate = blue, tropical = red) and shape denoting species. Thin, light species-level LOESS smooths (with shaded 95% confidence intervals) are overlaid by thicker biogeography-level (i.e., temperate and tropical) LOESS smooths (with shaded 95% confidence intervals).

Under well-watered conditions and low VPD, temperate species showed a breaking point (BP) at 35°C (Davies *p* = 0.0203), with A_net_ increasing significantly until this threshold (*R²* = 0.18, *p* = 0.0041, slope = 0.032) and subsequently decreasing, however non-significantly, above it (*R²* = 0.08, *p* = 0.13, slope = –0.054; Fig. S1a). By contrast, tropical species displayed no BP (Davies *p* = 0.132), and A_net_ continuously increased up to 40°C (*R²* = 0.38, *p* = 5.2 × 10⁻⁸, slope = 0.81; Fig. S1b). When T_air_ and VPD increased simultaneously, no BP was detected in either group (Davies *p* = 0.629 and *p* = 0.162, respectively). A_net_ of temperate species showed no significant response, whereas A_net_ of tropical species still exhibited a significant positive trend (*R²* = 0.12, *p* = 0.038, slope = 0.047; Figs. S1c, d). At constant 35°C, A_net_ of temperate species showed no significant response (Fig. S1f), while tropical species exhibited a significant decrease of A_net_ with rising VPD (Davies *p* = 0.492, *R²* = 0.13, *p* = 0.003, slope = –0.099; Fig. S1e).

Under drought, responses shifted (Fig. S2). In temperate species, A_net_ decreased steadily with increasing T_air_ (Davies *p* = 0.199, *R²* = 0.07, *p* = 0.02, slope = –0.022; Fig. S2a). However, in tropical species, a BP occurred at 35°C (Davies *p* = 0.0021), where A_net_ increased significantly until the BP (*R²* = 0.21, *p* = 0.0034, slope = 0.08), but declined significantly above it (*R²* = 0.19, *p* = 0.025, slope = –0.202; Fig. S2b). With increasing T_air_ and VPD, temperate species showed no significant response (Fig. S2c), whereas tropical species decreased continuously (Davies *p* = 0.64, *R²* = 0.17, *p* = 6.4 × 10⁻⁴, slope = –0.029; Fig. S2d). At a stable T_air_ of 35°C, A_net_ decreased significantly with rising VPD in both groups (*R²* = 0.23, *p* = 5.9 × 10⁻⁵, slope = –0.184; Fig. S2e in temperate and *R²* = 0.19, *p* = 9 × 10⁻⁵, slope = –0.151; Fig. S2f in tropical species).

Overall, temperate species under well-watered conditions benefited from increasing T_air_ only until 35°C when VPD remained low (Fig. S1a), but not when VPD simultaneously increased (Fig. S1c) or when drought-exposed (Fig. S2a, c). Their A_net_ was unresponsive to rising VPD at 35°C when well-watered (Fig. S1e), but became VPD sensitive when drought-exposed (Fig. S1e). A_net_ of tropical species, on the other hand, benefited more from increasing T_air_, even during soil drought, when VPD remained low (Figs. S1b, S2b), but only when well-watered when T_air_ and VPD increased simultaneously (Figs. S1d, S2d), and not when VPD increased at a stable T_air_, independent of soil moisture availability (Figs. S1f, S2f).

### T_air_ and VPD response of stomatal conductance in light- and dark-adapted leaves

Responses of stomatal conductance in the light (g_s_) partly mirrored those of photosynthesis but often revealed an independent response of the two processes, again with distinct differences between temperate and tropical species (Fig. 2; Figs. S3, S4).

**Figure 2:**
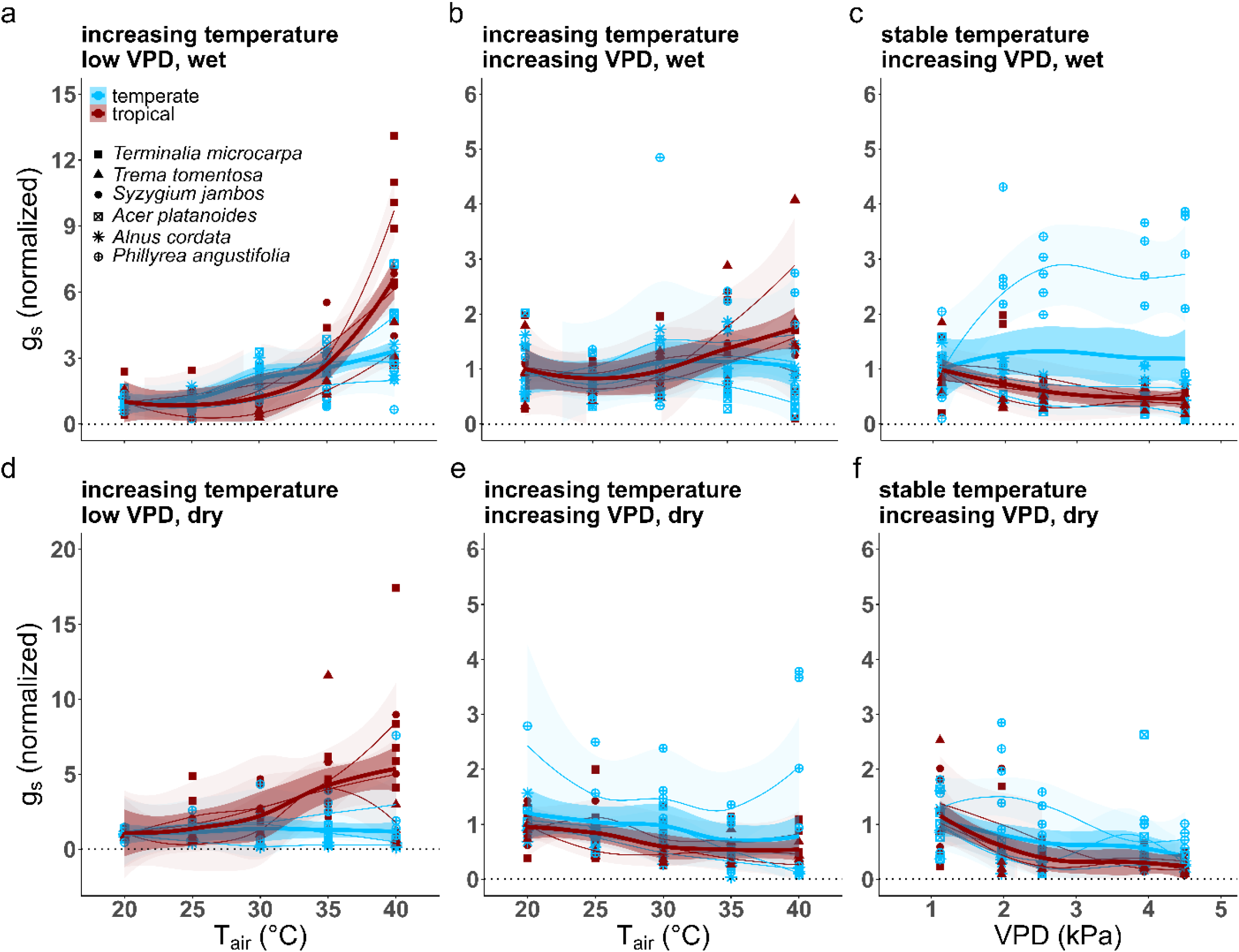
Response of the normalized stomatal conductance (g_s_, normalized to the species-specific average value at day 2) when temperature (T_air_) increases while vapour pressure deficit (VPD) remains low (a, d), when T_air_ and VPD simultaneously increase (b, e), and when VPD increases at a stable T_air_ of 35°C (c, f) of well-watered plants (a, b, c) or drought-exposed (d, e, f) plants. Points show individual measurements, with color indicating biogeographic origin (temperate = blue, tropical = red) and shape denoting species. Thin, light species-level LOESS smooths (with shaded 95% confidence intervals) are overlaid by thicker biogeography-level (i.e., temperate and tropical) LOESS smooths (with shaded 95% confidence intervals).

**Figure 3:**
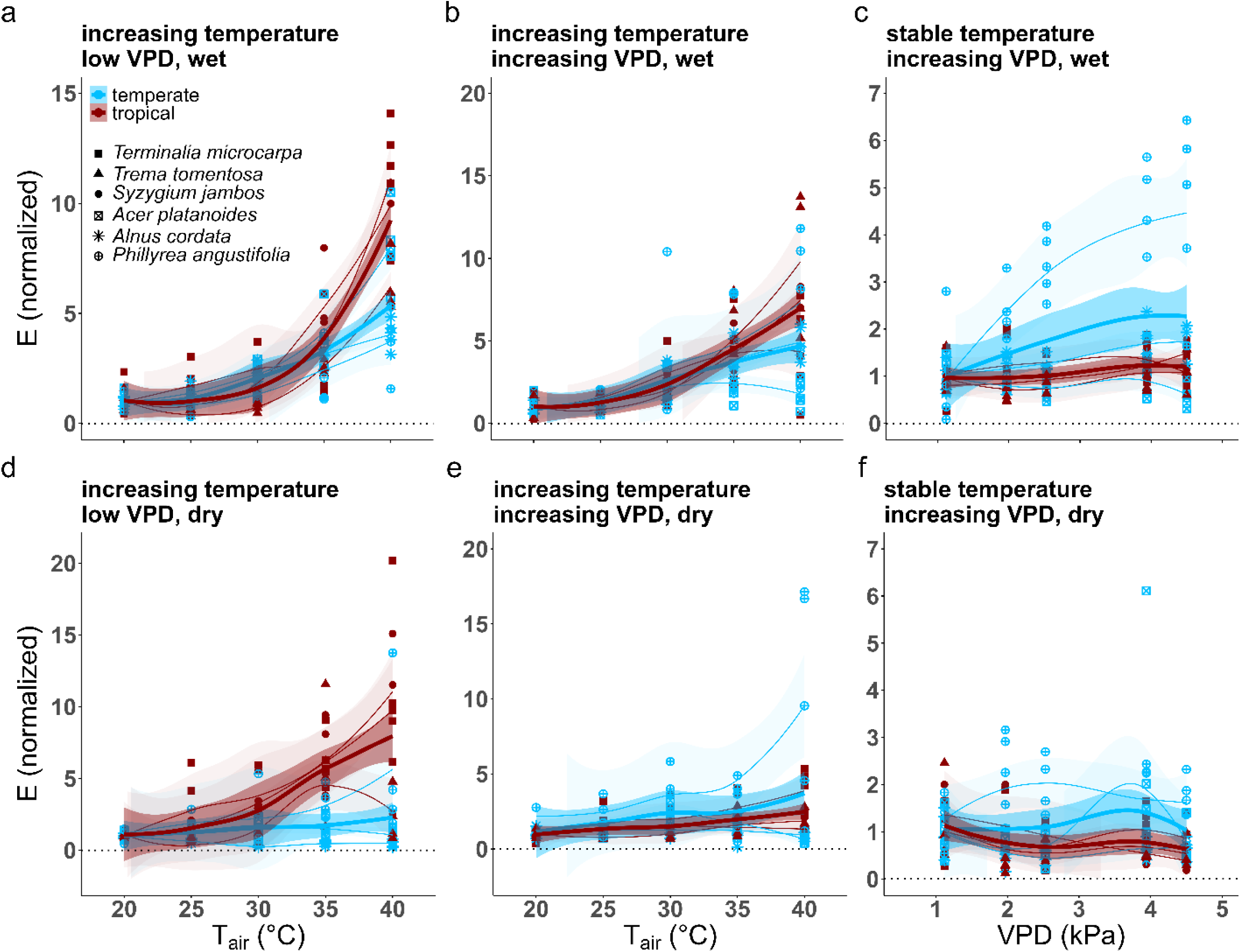
Response of the normalized transpiration (E, normalized to the species-specific average value at day 2) when temperature (T_air_) increases while vapour pressure deficit (VPD) remains low (a, d), when T_air_ and VPD simultaneously increase (b, e), and when VPD increases at a stable T_air_ of 35°C (c, f) of well-watered plants (a, b, c) or drought-exposed (d, e, f) plants. Points show individual measurements, with color indicating biogeographic origin (temperate = blue, tropical = red) and shape denoting species. Thin, light species-level LOESS smooths (with shaded 95% confidence intervals) are overlaid by thicker biogeography-level (i.e., temperate and tropical) LOESS smooths (with shaded 95% confidence intervals).

Under well-watered and low VPD conditions, temperate species exhibited a continuous increase of g_s_ with increasing T_air_ (no BP, Davies *p* = 0.0876; *R²* = 0.43, *p* = 2.7 × 10⁻¹⁰, slope = 0.119; Fig. S3a). In tropical species, a BP was detected at 33.65°C (Davies *p* = 8.3 × 10⁻⁷), with stable g_s_ below this threshold (*R²* = 0.01, *p* = 0.58, slope = 0.014) and a strong increase thereafter (*R²* = 0.44, *p* = 2.2 × 10⁻⁴, slope = 0.87; Fig. S3b). When T_air_ and VPD rose simultaneously, a significant increase in g_s_ occurred only in tropical species (*R^2^* = 0.15, *p* = 0.0015, slope = 0.04; Figs. S3c, d). At constant 35°C, temperate species showed no significant VPD effect (Fig. S3e), while tropical species exhibited a strong continuous decline in g_s_ (*R²* = 0.26, *p* = 2.6 × 10⁻⁵, slope = –0.163; Fig. S3f).

When drought-exposed, temperate species showed no significant changes in g_s_ with increasing T_air_ (Fig. S4a), while tropical species displayed a continuous increase (*R²* = 0.26, *p* = 1.6 × 10⁻⁵, slope = 0.219; Fig. S4b). When T_air_ and VPD increased together, g_s_ of temperate species remained unresponsive (Fig. S4c), whereas g_s_ of tropical species steadily declined (*R²* = 0.17, *p* = 6 × 10⁻⁴, slope = –0.026; Fig. S4d). At constant 35°C, g_s_ steadily decreased with rising VPD in both groups (*R²* = 0.11, *p* = 0.0037, slope = –0.159; Fig. S4e in temperate, and *R²* = 0.20, *p* = 2.2 × 10⁻⁴, slope = –0.189; Fig. S4f in tropical species).

Responses of stomatal conductance of dark-adapted leaves (g_s dark_) to T_air_ and VPD showed similar overall trends as during the light. However, the general sensitivity of g_s dark_ differed from the observed response in the light, and between temperate and tropical species (Figs. S5–S7).

Under well-watered conditions, temperate species exhibited a BP at 34.9°C (Davies *p* = 0.0122). Below this threshold, g_s dark_ increased only slightly with rising T_air_ (*R²* = 0.13, *p* = 0.017, slope = 0.165), above the BP, the increase in g_s_ was stronger (*R²* = 0.21, *p* = 0.013, slope = 0.926; Fig. S6a). Similar to the light-adapted leaf, g_s_ of tropical species displayed a BP at 32.7°C (Davies *p* = 0.00145), with no significant change below it (*R²* = 0.01, *p* = 0.64, slope = 0.013) and a strong increase above (*R²* = 0.20, *p* = 0.021, slope = 0.63; Fig. S6b). When T_air_ and VPD increased simultaneously, g_s dark_ remained unchanged in both groups (Fig. S6c, d). At constant 35°C, temperate and tropical species exhibited a significant constant decline of g_s dark_ with increasing VPD (*R²* = 0.07, *p* = 0.025, slope = –0.144; Fig. S6e, and *R²* = 0.30, *p* = 3.7 × 10⁻⁶, slope = –0.233; Fig. S6f, respectively).

Under drought, g_s dark_ of temperate and tropical species increased with rising T_air_ (*R²* = 0.06, *p* = 0.047, slope = 0.032; Fig. S7a and *R²* = 0.14, *p* = 0.0026, slope = 0.127; Fig. S7b, respectively). When T_air_ and VPD rose simultaneously, temperate species exhibited a significant constant decline in g_s dark_ (*R²* = 0.29, *p* = 1.2 × 10⁻⁶, slope = –0.036; Fig. S7c), whereas tropical species showed no significant change (Fig. S7d). At constant 35°C, g_s dark_ of temperate and tropical species displayed a BP at 2.31 kPa VPD (Davies *p* = 0.00113) and 2.53 kPa (Davies p = 9.8 × 10⁻^4^), respectively. Below this threshold, g_s dark_ decreased strongly (*R²* = 0.19, *p* = 0.016, slope = –0.559Fig. S7e and *R²* = 0.20, *p* = 0.034, slope = –0.588, Fig. S7e, respectively), while no further decrease was observed beyond it (*R²* = 0.00, *p* = 0.89, slope = –0.004; Fig. S7e and *R²* = 0.02, *p* = 0.42, slope = –0.020; Fig. S7f, respectively).

The ratio of stomatal conductance in the light to that in the dark (g_s_-ratio, g_s_ divided by g_s dark_) revealed additional contrasts between temperate and tropical species and between water treatments (Figs. S8–S10), further revealing different responses of stomatal regulation in light- and dark-adapted leaves.

Under wet conditions and low VPD, temperate species showed exhibited a significant steady decline in the ratio from 3.36 ± 0.69 (SE) at 20°C to 1.30 ± 0.14 at 40°C (*R²* = 0.16, *p* = 0.00071, slope = –0.107; Fig. S9a), meaning that g_s_ was initially substantially higher in the light than in the dark, but this difference vanished and they became very similar at 40°C. In tropical species, no significant response occurred (Fig. S9b), meaning that g_s_ in light- and dark-adapted leaves increased to a similar extent. When T_air_ and VPD increased simultaneously, the ratio of neither group showed significant responses (Figs. S9c, d). At constant 35°C, temperate species exhibited a BP at 1.97 kPa (Davies *p* = 0.0456). Before this threshold, the g_s_-ratio increased steeply from 1.97 ± 0.32 to 4.27 ± 0.88 (*R²* = 0.18, *p* = 0.021, slope = 2.707), whereas above it no significant VPD response was detected (*R²* = 0.00, *p* = 0.94, slope = –0.025; Fig. S9e). Tropical species, on the other hand, had a BP at a higher VPD of 3.94 kPa (Davies *p* = 0.0453). Below the BP, the g_s_- ratio showed no significant trend (*R²* = 0.09, *p* = 0.086, slope = 0.664), but above it, the g_s_-ratio increased strongly (*R²* = 0.19, *p* = 0.029, slope = 4.722; Fig. S9f), showing a much stronger reduction of g_s dark_ and gs during the light at the highest VPD. Overall, this shows a different regulation of stomatal conductance in the light and the dark in response to increasing T_air_ and VPD among temperate species at a low VPD (Fig. S9a), but not if they increase simultaneously (Fig. S9c). The response of g_s_ and g_s dark_ in tropical species, however, responded very similarly in all but the most extreme VPD conditions at a stable T_air_ of 35°C.

Under drought and a low VPD, the g_s_-ratio of temperate species displayed a BP at 25.65°C (Davies *p* = 0.0307), with an increase between 20°C to 25°C (*R²* = 0.09, *p* = 0.12, slope = 0.26), and a significant steady decrease until 40°C (*R²* = 0.12, *p* = 0.027, slope = –0.184; Fig. S10a), driven by the steady increase of g_s dark_ (Fig. S7a) while g_s_ did not change (Fig. S4a). In tropical species, the g_s_-ratio increased steadily with T_air_ (*R²* = 0.12, *p* = 0.0057, slope = 0.071; Fig. S10b), caused by the stronger increase of g_s_ in the light (Fig. S4b) than in the dark (Fig. S7b). With simultaneous increases in Tair and VPD, the g_s_-ratio of neither temperate nor tropical species showed significant responses (Figs. S10c, d). At constant 35°C, the g_s_-ratio of temperate species showed no significant response to rising VPD (Fig. S10e), whereas tropical species exhibited a steady increase (*R²* = 0.08, *p* = 0.0024, slope = 0.395, Fig. S10f).

### T_air_ and VPD response of transpiration in light- and dark-adapted leaves

Unlike g_s_, transpiration (E) showed a much more consistent increase with T_air_ under low and rising VPD, rather constant with rising VPD at a constant T_air_, and less differentiated response between temperate and tropical species within watering regimes (Figs. S11, S12).

Under well-watered conditions and low VPD, temperate species exhibited a BP at 31.6°C (Davies *p* = 0.00114). Below this threshold, E increased slightly (*R²* = 0.24, *p* = 0.0008, slope = 0.077), while above it the increase was much stronger (*R²* = 0.24, *p* = 0.006, slope = 0.41; Fig. S11a). Tropical species showed a BP at 33.5°C (Davies *p* = 2.7 × 10⁻¹⁰). Below this threshold, E did not respond to rising T_air_ (*R²* = 0.07, *p* = 0.11, slope = 0.05), while above the BP, E increased sharply (*R²* = 0.61, *p* = 2.3 × 10⁻⁶, slope = 1.169; Fig. S11b). When T_air_ and VPD increased simultaneously, E of temperate species rose steadily (*R²* = 0.31, *p* = 3.5 × 10⁻⁷, slope = 0.200; Fig. S11c), while tropical species exhibited a BP at 28.2°C (Davies *p* = 0.0292). Below this threshold, E did not respond significantly (*R²* = 0.13, *p* = 0.073, slope = 0.070), while we observed a strong increase above (*R²* = 0.38, *p* = 3.3 × 10⁻⁵, slope = 0.457; Fig. S11d). At constant 35°C, temperate species showed a significant steady increase with VPD (*p* = 0.0021, slope = 0.394; Fig. S11e), while E of tropical species exhibited no significant response (Fig. S11f).

Under drought, E of temperate species at a constant low VPD did not respond significantly to rising T_air_ (Fig. S12a), while it slightly increased under simultaneous increasing VPD (*R²* = 0.09, *p* = 0.0084, slope = 0.124, Fig. S12c). In tropical species, E increased significantly with T_air_, both under low (*R²* = 0.37, *p* = 8.8 × 10⁻⁸, slope = 0.342; Fig. S12b) and simultaneous increasing VPD (*R²* = 0.20, *p* = 2.3 × 10⁻⁴, slope = 0.342; Fig. S12d). At constant 35°C, E of neither group showed significant VPD responses (Figs. S12e, f).

Overall, this demonstrates the importance of the interplay between g_s_ and VPD in regulating E, with g_s_ needing to be upregulated when T_air_ increases to increase E when VPD remains low, but can remain unchanged when VPD simultaneously rises.

Similar to light-adapted leaves, transpiration of dark-adapted leaves (E_dark_) increased with T_air_ in both temperate and tropical species, with distinct BPs under well-watered conditions and low VPD but largely linear responses under increasing VPD or when drought-exposed (Figs. S13-S15).

Under wet conditions and low VPD, temperate species showed a BP at 33.9°C (Davies *p* = 0.00104). Below this threshold, E_dark_ increased modestly (*R²* = 0.12, *p* = 0.018, slope = 0.137), while above it the increase was much steeper (*R²* = 0.24, *p* = 0.0076, slope = 1.70; Fig. S14a). Tropical species exhibited a BP at 32.5°C (Davies *p* = 0.00232). Below this point, no response to T_air_ occurred (*R²* = 0.00, *p* = 0.97, slope = 0.001), but above it, E_dark_ increased strongly (*R²* = 0.19, *p* = 0.027, slope = 0.932; Fig. S14b). When T_air_ and VPD increased simultaneously, E_dark_ of temperate and tropical species showed significant steady increases (*R²* = 0.25, *p* = 6.4 × 10⁻⁶, slope = 0.304; and *R²* = 0.26, *p* = 1.5 × 10⁻⁵, slope = 0.394; respectively, Fig. S14c, d). At constant 35°C, temperate species showed no VPD response (Fig. S14e), whereas tropical species exhibited a significant decrease of E_dark_ with increasing VPD (*R²* = 0.07, *p* = 0.037, slope = –0.126; Fig. S14f).

Under drought, E_dark_ of temperate and tropical species increased steadily (*R²* = 0.16, *p* = 0.00049, slope = 0.098; Fig. S15a, and *R²* = 0.21, *p* = 1.3 × 10⁻⁴, slope = 0.244; Fig. S15b, respectively). With simultaneous increases in T_air_ and VPD, only temperate species showed a significant steady increase (*R²* = 0.14, *p* = 0.0012, slope = 0.053; Fig. S15c, d). At constant 35°C, temperate species exhibited a BP at 2.4 kPa (Davies *p* = 0.0431). Below this threshold, E_dark_ decreased slightly but not significantly (*R²* = 0.06, *p* = 0.18, slope = –0.332), whereas above it a significant increase occurred (*R²* = 0.09, *p* = 0.046, slope = 0.135; Fig. S15e). In tropical species, E_dark_ decreased steadily in response to rising VPD (*R²* = 0.17, *p* = 0.00091, slope = –0.176; Fig. S15f). This indicates that E of dark-adapted leaves is more conservatively regulated than transpiration in light-adapted leaves at high T_air_ when water availability is restricted, but still remain responsive to changes in T_air_ and VPD.

The ratio of transpiration in the light to that in the dark (E-ratio; E divided by E_dark_) showed less response compared to absolute fluxes, further verifying mostly similar responses of E independent of light availability (Figs. S16-18).

However, under wet conditions and low VPD, temperate species exhibited a significant steady decrease in the ratio with increasing T_air_ (*R²* = 0.16, *p* = 0.00057, slope = –0.128; Fig. S17a), showing that the E and E_dark_ become more and more similar. No significant change of the E-ratio was observed in tropical species when T_air_ increased, both at a low VPD (Fig. S17b) and increasing VPD (Fig. S17d), and in temperate species when T_air_ and VPD increased simultaneously (Fig. S17c) or VPD increased at constant 35°C (Fig. S17e). In contrast, the E-ratio of tropical species exhibited a strong linear increase with rising VPD at 35°C (*R²* = 0.21, *p* = 0.00027, slope = 0.902; Fig. S17f), showing their stronger VPD sensitivity of E_dark_ compared to E.

Under drought, when T_air_ increased while VPD remained low, temperate species displayed a BP at 25°C (Davies *p* = 0.033). Below this point, the E-ratio did increase but not overall significantly, whereas above it the E-ratio decreased strongly, but again not significantly (Fig. S18a). The E- ratio of tropical species increased steadily with rising T_air_ (*R²* = 0.11, *p* = 0.011, slope = 0.081; Fig. S18b). When T_air_ and VPD increased simultaneously or VPD increased at a constant 35°C, the E- ratio of neither group showed a significant response (Figs. S18c, d, e, f).

These findings indicate that E, both in the light and the dark, is mostly regulated in a similar range in response to rising T_air_, especially when water is not limited, while their ratio remains largely constant in response to rising VPD at a constant T_air_.

### T_air_ and VPD response of the intrinsic water use efficiency

The intrinsic water use efficiency (WUE_i_, A_net_ divided by g_s_) showed contrasting T_air_ and VPD responses between biogeographic groups and water regimes, with distinct BPs under both wet and dry conditions (Fig. 4; Figs. S19, S20).

**Figure 4:**
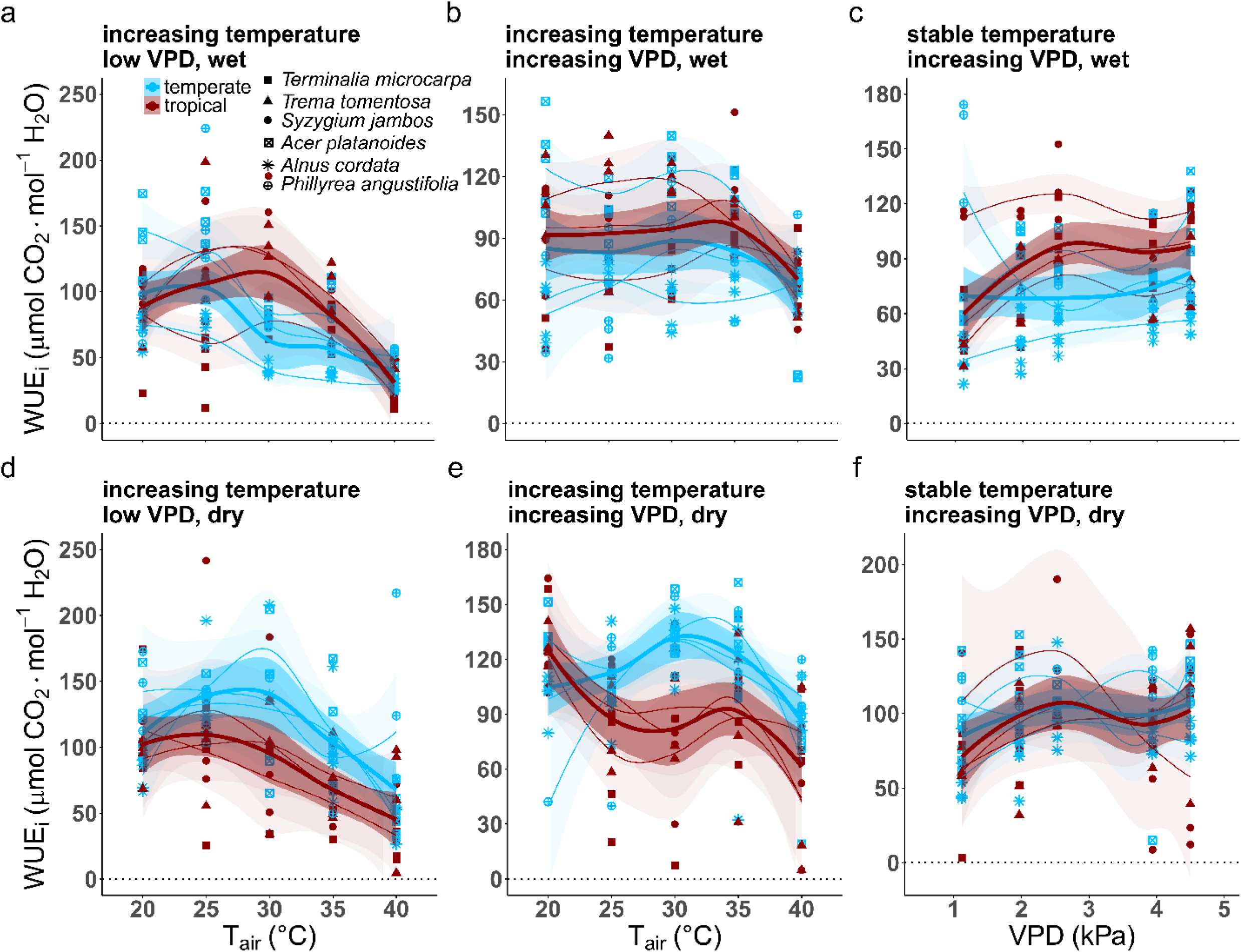
Response of the intrinsic water use efficiency (WUE_i_ (A_net_ divided by g_s_), when Tair increases while vapour pressure deficit (VPD) remains low (a, d), when Tair and VPD simultaneously increase (b, e), and when VPD increases at a stable Tair of 35°C (c, f) of well-watered plants (a, b, c) or drought-exposed (d, e, f) plants. Points show individual measurements, with color indicating biogeographic origin (temperate = blue, tropical = red) and shape denoting species. Thin, light species-level LOESS smooths (with shaded 95% confidence intervals) are overlaid by thicker biogeography-level (i.e., temperate and tropical) LOESS smooths (with shaded 95% confidence intervals).

Under well-watered conditions and low VPD, WUE_i_ of temperate species exhibited a BP at 25°C (Davies *p* = 0.00878). Below this threshold, WUE_i_ remained stable (*R²* = 0.11, *p* = 0.079, slope = 5.568), while above it WUE_i_ decreased significantly with increasing T_air_ (*R²* = 0.20, *p* = 0.002, slope = –4.185; Fig. S19a). In tropical species, a BP occurred at 31.9°C (Davies *p* = 0.00226), with no significant response below (*R²* = 0.06, *p* = 0.13, slope = 3.645), but with WUE_i_ declining sharply above (*R²* = 0.28, *p* = 0.005, slope = –12.120; Fig. S19b). When T_air_ and VPD increased simultaneously, WUE_i_ remained unchanged in temperate species (Fig. S19c), but exhibited a BP at 35°C in tropical species (Davies *p* = 0.00649). Below the BP, WUE_i_ remained constant (*R²* = 0.00, *p* = 0.76, slope = 0.331), while above it, WUE_i_ decreased steeply (*R²* = 0.50, *p* = 7.1 × 10⁻⁵, slope = –6.715; Fig. S19d). At constant 35°C, WUE_i_ remained stable in temperate species (Fig. S19e), whereas tropical species showed a steady increase with rising VPD (*R²* = 0.14, *p* = 0.0024, slope = 8.277; Fig. S19f).

Under drought, WUE_i_ responses to T_air_ further differentiated among temperate and tropical species (Fig. S20). Temperate species showed a BP with low VPD at 30°C (Davies *p* = 0.00354): below this point, WUE_i_ did not change (*R²* = 0.04, *p* = 0.32, slope = 3.101), but above it declined markedly (*R²* = 0.29, *p* = 2.4 × 10⁻⁴, slope = –8.605; Fig. S20a). With simultaneously rising VPD, they showed a BP at 33.14°C (Davies *p* = 0.000706). Below the BP, a slight but significant increase was observed (*R²* = 0.10, *p* = 0.039, slope = 2.05), while above it, WUE_i_ decreased strongly (*R²* = 0.27, *p* = 0.0036, slope = –6.619; Fig. S20c). On the other hand, WUE_i_ of tropical species decreased continuously with T_air_ independent of VPD (*R²* = 0.20, *p* = 2.1 × 10⁻⁴, slope = –2.764; Fig. S20b and *R²* = 0.12, *p* = 0.0061, slope = –1.935; Fig. S20d, respectively). At constant T_air_, WUE_i_ of neither group showed significant VPD responses (Fig. S20e, f).

### T_air_ and VPD response of the water use efficiency

Unlike WUE_i_, the actual water-use efficiency (WUE, A_net_ divided by E) declined more consistently with rising T_air_, independent of VPD and soil moisture availability (Fig. 5). We found distinct breaking points (BPs) in response to increasing T_air_ under a low and rising VPD, both in temperate and tropical species, and a consistent decrease in response to rising VPD at a constant 35°C (Figs. S21, S22).

**Figure 5:**
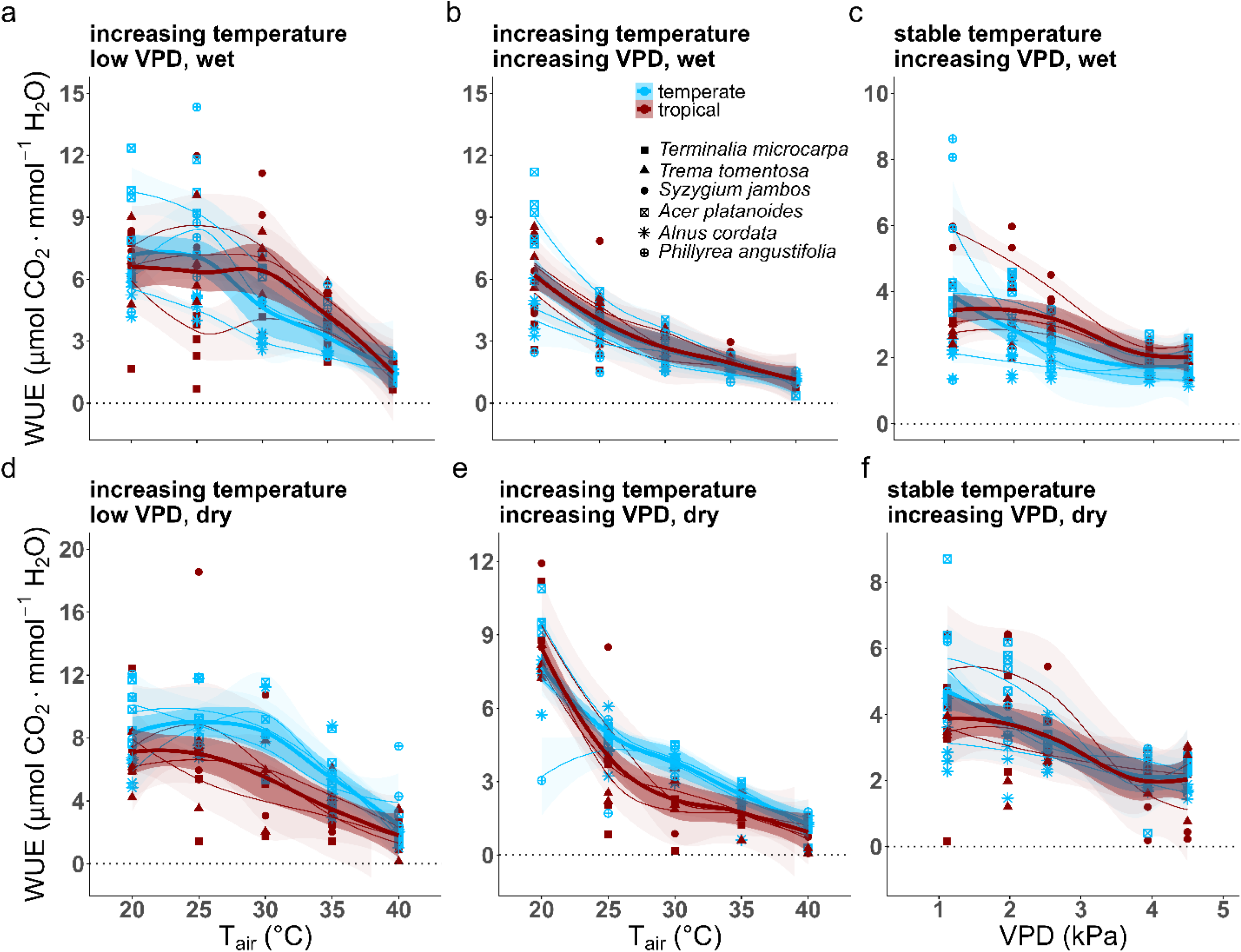
Response of the normalized water use efficiency (WUE (A_net_ divided by E), normalized to the species-specific average value at day 2) when Tair increases while vapour pressure deficit (VPD) remains low (a, d), when Tair and VPD simultaneously increase (b, e), and when VPD increases at a stable Tair of 35°C (c, f) of well-watered plants (a, b, c) or drought-exposed (d, e, f) plants. Points show individual measurements, with color indicating biogeographic origin (temperate = blue, tropical = red) and shape denoting species. Thin, light species-level LOESS smooths (with shaded 95% confidence intervals) are overlaid by thicker biogeography-level (i.e., temperate and tropical) LOESS smooths (with shaded 95% confidence intervals).

Under well-watered conditions and low VPD, temperate species showed a BP at 25°C (Davies *p* = 0.00291). Below this threshold, WUE remained constant (*R²* = 0.03, *p* = 0.33, slope = 0.199), while above it WUE declined sharply (*R²* = 0.35, *p* = 2.15 × 10⁻⁵, slope = –0.409; Fig. S21a). In tropical species, a BP occurred at 30.8°C (Davies *p* = 0.02), where WUE did not respond below (*R²* = 0.00, *p* = 0.78, slope = 0.039), but above it a strong decline was observed (*R²* = 0.40, *p* = 5.3 × 10⁻⁴, slope = –0.564; Fig. S21b). When T_air_ and VPD increased simultaneously, temperate species showed a BP at 25.6°C (Davies *p* = 0.003). Before this point, WUE decreased strongly (*R²* = 0.33, *p* = 0.002, slope = –0.549), and after it continued to decline more gradually (*R²* = 0.57, *p* = 3.8 × 10⁻⁹, slope = –0.158; Fig. S21c). Similarly, WUE of tropical species displayed a BP at 26.7°C (Davies *p* = 0.0193). Below this threshold, WUE decreased more steeply (*R²* = 0.30, *p* = 0.0039, slope = –0.435), while above it the decline was less pronounced (*R²* = 0.67, *p* = 2.9 × 10⁻¹⁰, slope = –0.159; Fig. S21d). At constant T_air_, temperate and tropical species showed steady decreases with rising VPD (*R²* = 0.25, *p* = 5.9 × 10⁻⁶, slope = –0.573 and *R²* = 0.37, *p* = 1.2 × 10⁻⁷, slope = – 0.527, respectively; Fig. S21e, f).

Under drought, the pattern remained largely remained (Fig. S22). Temperate species showed a BP at 29.9°C (Davies *p* = 0.00158). Below this T_air_, WUE remained unchanged (*R²* = 0.00, *p* = 0.95, slope = 0.015), whereas above it, a pronounced decrease occurred (Fig. S22a). In tropical species, WUE decreased steadily (*R²* = 0.36, *p* = 2.5 × 10⁻⁷, slope = –0.264; Fig. S22b). When both T_air_ and VPD increased, temperate species exhibited a BP at 25°C (Davies *p* = 2.1 × 10⁻⁵). Below this point, WUE declined steeply (*R²* = 0.57, *p* = 5.9 × 10⁻⁶, slope = –0.710), whereas above it the decline was weaker (*R²* = 0.83, *p* = 5.6 × 10⁻¹⁸, slope = –0.252; Fig. S22c). Similar again in tropical species, a BP was detected at 26.1°C (Davies *p* = 0.0193). Before this threshold, WUE decreased strongly (*R²* = 0.50, *p* = 7.2 × 10⁻⁵, slope = –0.831), and after it the decline flattened (*R²* = 0.32, *p* = 3.8 × 10⁻⁴, slope = –0.135; Fig. S22d). At constant 35°C, both temperate and tropical species showed steady decreases in WUE with rising VPD (*R²* = 0.46, *p* = 3.1 × 10⁻¹¹, slope = –0.762 and *R²* = 0.22, *p* = 1.2 × 10⁻⁴, slope = –1.310, respectively; Figs. S22e, f).

### The relationship between A_net_ and g_s_ in response to rising T_air_ and VPD

Across all treatments, A_net_ and g_s_ were strongly correlated, but the strength and slope of their relationship varied with T_air_, VPD, and biogeographic group (Fig. 6; Fig. S23).

**Figure 6:**
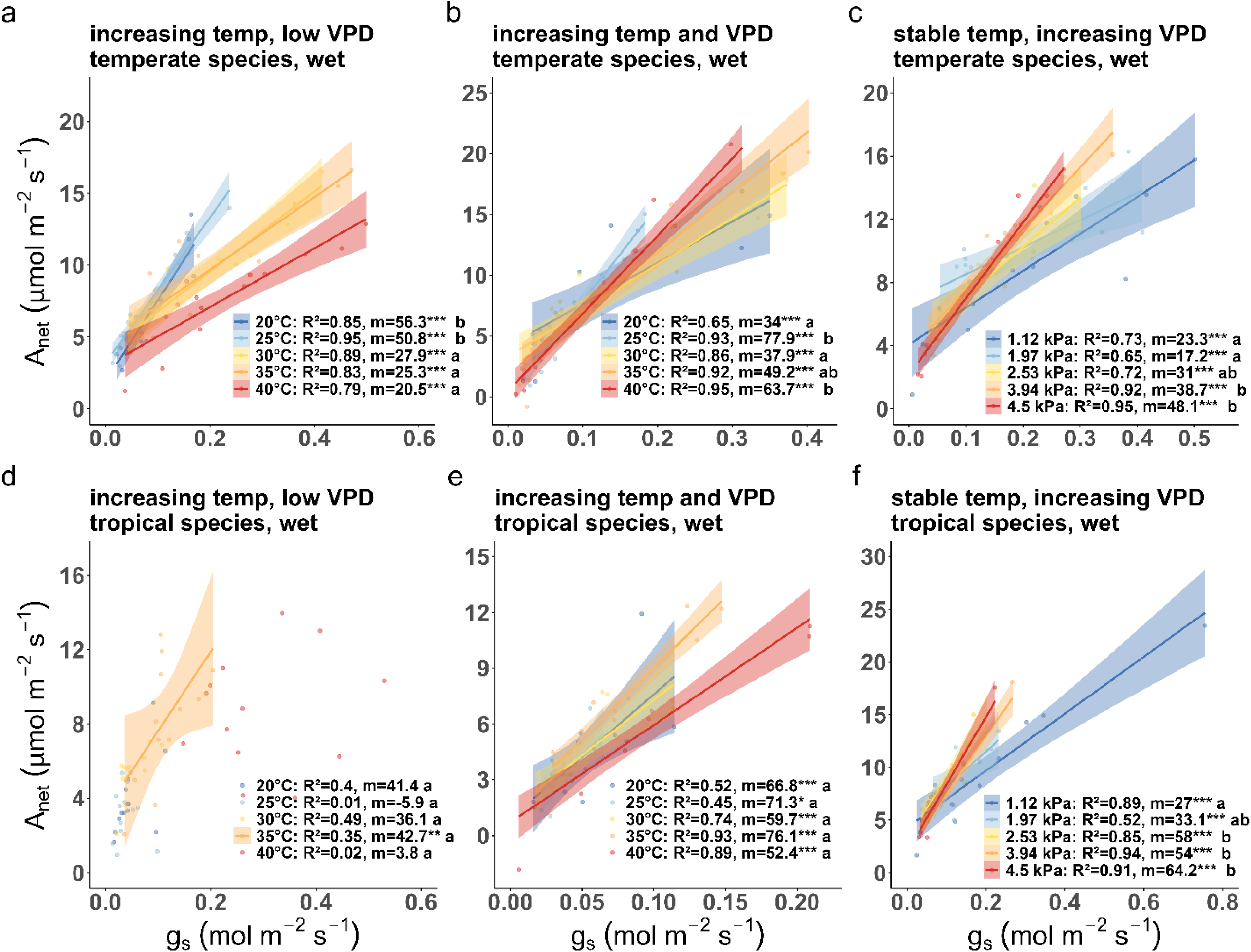
Changes of the relationship between net assimilation (A_net_) and stomatal conductance (g_s_) when temperature (T_air_) increases while vapour pressure deficit (VPD) remains low (a, d), Tair and VPD are simultaneously increasing (b, e), and VPD increases at a stable T_air_ of 35°C (c, f) in temperate (a, b, c) and tropical (d, e, f) tree species of well-watered plants. Points show individual measurements; colored lines are linear fits plotted only when slopes are significant (p < 0.05) from global interaction models (A_net_ ∼ g_s_ × Temp or A_net_ ∼ g_s_ × VPD), with shaded 95% confidence intervals. Legends report per-level R² and slope m (A_net_/g_s_) with significance asterisks (* p < 0.05, ** p < 0.01, *** p < 0.001), and compact letter displays summarize pairwise slope differences.

Under well-watered conditions, temperate species showed consistently high correlations between A_net_ and g_s_ (Fig. 6a–c). At low VPD, *R²* values ranged from 0.79 to 0.95 across temperatures. Slopes (*m*) declined from 56.3 (b) at 20°C (*R²* = 0.85***) to 20.5 (a) at 40°C (*R²* = 0.79***), indicating a progressive reduction in photosynthetic gain per unit stomatal conductance. When Tair and VPD increased simultaneously, slopes varied from 34 (a) at 20°C (*R²* = 0.65***) to 63.7 (b) at 40°C (*R²* = 0.95***; Fig. 6b), but no clear trend in their relationship was visible. At constant 35°C and rising VPD, the A_net_–g_s_ relationship steepened, with *m* = 23.3 (a) at 1.12 kPa (*R²* = 0.73***) and *m* = 48.1 (b) at 4.5 kPa (*R²* = 0.95***; Fig. 6c).

In tropical species, coupling between A_net_ and g_s_ was weaker and less consistent under wet conditions when VPD was low (Fig. 6d), where regressions were only significant at 35°C (*R²* = 0.35, *m* = 42.7***). When Tair and VPD rose together, slopes remained relatively constant and did not significantly differ from each other (*m* = 66.8 (a) to 76.1 (a); *R²* = 0.45***–0.93***; Fig. 6e). At stable Tair, increasing VPD produced a progressive steepening of the A_net_-g_s_ relationship: *m* = 27 (a) at 1.12 kPa (*R²* = 0.89***) to *m* = 64.2 (b) at 4.5 kPa (*R²* = 0.91***; Fig. 6f).

Under drought, A_net_–g_s_ coupling was more consistent, with mostly no significant variation, though slope magnitudes differed (Fig. S23). In temperate species, slopes were relatively stable across Tair treatments at a constant low VPD (*m* = 66 (a) to 84.8 (a); *R²* = 0.68***–0.82***; Fig. S23a) and increased when T_air_ and VPD rose simultaneously (*m* = 98.4 (a) at 20°C to 125 (a) at 35°C; *R²* = 0.48***–0.99***; Fig. S23b). Under constant T_air_, *m* rose from 44 (a) at 1.12 kPa (*R²* = 0.84***) to 75.8 (a) at 4.5 kPa (*R²* = 0.80***; Fig. S23c).

In tropical species, when drought-exposed under constant low VPD, A_net_–g_s_ correlations remained absent at 20–30°C but became significant at higher T_air_ (*R²* = 0.78***, *m* = 39.3 (b) at 35°C; *R²* = 0.43**, *m* = 13.9 (a) at 40°C; Fig. S23d). When T_air_ and VPD increased together, slopes remained steep and uniform (*m* = 83.4 (a) to 113.3 (a); *R²* = 0.72***–0.89***; Fig. S23e). At constant 35°C, rising VPD further strengthened the A_net_–g_s_ coupling, but the regressions did not significantly differ from each other: *m* = 68.4 (a) at 1.12 kPa (*R²* = 0.89***) to *m* = 132.3 (a) at 4.5 kPa (*R²* = 0.87***; Fig. S23f).

Overall, the A_net_–g_s_ relationship weakened with increasing T_air_ when VPD remained low in temperate species, but remained strong under rising VPD and increased under drought exposure. In tropical species, coupling was largely absent under low VPD but strengthened under high VPD, particularly when soils were dry.

### The relationship between A_net_ and E in response to rising T_air_ and VPD

Coupling between A_net_ and E (slopes *m*) decreased with rising T_air_, both with a low and a simultaneously rising VPD, and was nearly absent in tropical species under low VPD (Fig. 7; Fig. S24).

**Figure 7:**
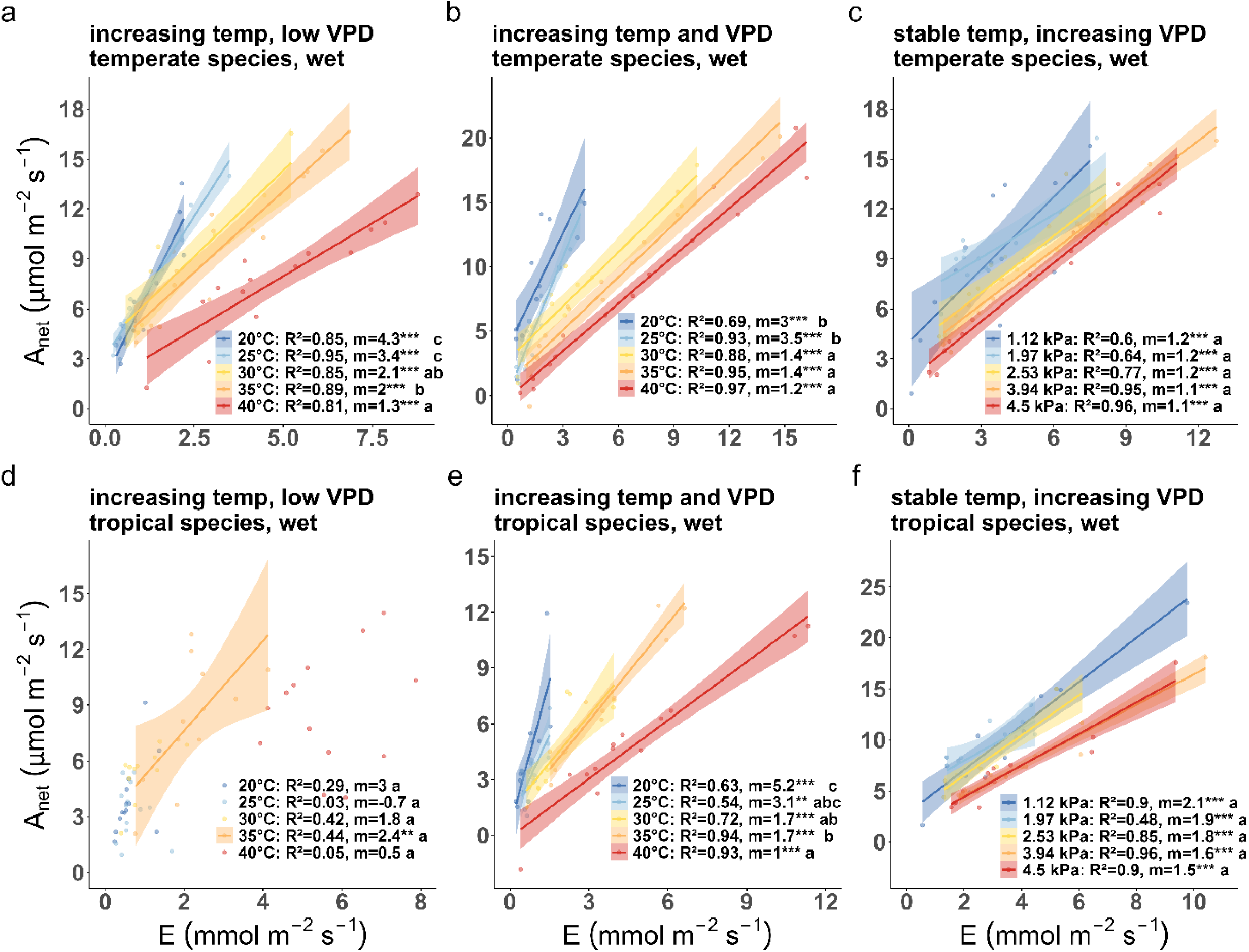
Changes of the relationship between net assimilation (A_net_) and transpiration (E) when temperature (T_air_) increases while vapour pressure deficit (VPD) remains low (a, d), Tair and VPD are simultaneously increasing (b, e), and VPD increases at a stable Tair of 35°C (c, f) in temperate (a, b, c) and tropical (d, e, f) tree species of well-watered plants. Points show individual measurements; colored lines are linear fits plotted only when slopes are significant (p < 0.05) from global interaction models (A_net_ ∼ g_s_ × Temp or A_net_ ∼ g_s_ × VPD), with shaded 95% confidence intervals. Legends report per-level R² and slope m (A_net_/g_s_) with significance asterisks (* p < 0.05, ** p < 0.01, *** p < 0.001), and compact letter displays summarize pairwise slope differences.

Under well-watered conditions, temperate species showed highly significant relationships at all T_air_ (Fig. 7a–c). At low VPD, *R²* ranged from 0.81–0.95, with slopes declining from 4.3 (c) at 20°C (*R²* = 0.85***) to 1.3 (a) at 40°C (*R²* = 0.81***), reflecting reduced photosynthetic gain per unit transpiration. When T_air_ and VPD rose simultaneously, relationships fell from 3.0 (b) at 20°C (*R²* = 0.69***) to 1.2 (a) at 40°C (*R²* = 0.97***; Fig. 7b). At constant 35°C, increasing VPD did not impact the A_net_-E relationship (*m* = 1.1–1.2 (a); *R²* = 0.60***–0.96***; Fig. 7c).

In tropical species, A_net_-E coupling was nearly absent at low VPD but strengthened under higher VPD (Fig. 7d–f). Relationships under rising T_air_ and low VPD were only significant at 35°C (*R²* = 0.44**, *m* = 2.4, Fig. 7d). When T_air_ and VPD increased together, *R²* values ranged 0.54***– 0.94*** and slopes decreased from 5.2 (c) at 20°C to 1.0 (a) at 40°C (Fig. 7e). At constant T_air_, increasing VPD did not significantly impact the relationships between Anet and E (1.5–2.1 (a), *R²* = 0.48***–0.96***; Fig. 7f).

Under drought, the general trends of the A_net_-E coupling remained unchanged (Fig. S24). In temperate species, slopes declined from 4.8 (b) at 20°C (*R²* = 0.78***) to 1.8 (a) at 40°C (*R²* = 0.77***; Fig. S24a). When T_air_ and VPD increased simultaneously, slopes decreased systematically from 7.5 (c) at 20°C (*R²* = 0.48***) to 1.5 (a) at 40°C (*R²* = 0.99***; Fig. S24b). At constant T_air_, increasing VPD yielded steady slopes of 1.6–2.6, which did not differ significantly ((a), *R²* = 0.49***–0.86***; Fig. S24c).

In drought-exposed tropical species, at constant low VPD and rising T_air_, no significant relationships occurred up to 30°C, but relationships became significant at 35°C (*R²* = 0.70***, *m* = 2.6 (b)) and 40°C (*R²* = 0.46**, *m* = 0.8 (a); Fig. S24d). When T_air_ and VPD rose together, relationships were consistent (*m* = 2.9–5.7 (a); *R²* = 0.55***–0.91***; Fig. S24e). At constant T_air_, the A_net_-E relationship weakened slightly with rising VPD, with *m* declining from 4.1*** (b) at 1.12 kPa (*R²* = 0.89) to 2.3*** (a) at 3.94 kPa (*R²* = 0.91; Fig. S24f).

Overall, A_net_-E coupling expressed a consistent decline with increasing T_air_, independent of VPD, in both climatic groups.

### The relationship between E and g_s_ in response to rising T_air_ and VPD

Under well-watered conditions, temperate species showed highly significant E*–*g_s_ coupling across the full T_air_ range (Fig. 8a–c). At low VPD, *R²* remained ≥ 0.97. Slopes increased slightly with T_air_ from 13*** (ab) at 20°C to 16.1*** (a) at 40°C (Fig. 8a). When both T_air_ and VPD increased, the relationship steepened strongly, with *m* rising from 11.7*** (a) at 20°C to 52.2*** (e) at 40°C (*R²* = 0.99–1; Fig. 8b). At constant T_air_, increasing VPD produced a similar pattern: *m* = 13.9*** (a) at 1.12 kPa (*R²* = 0.92) and *m* = 41.1*** (e) at 4.5 kPa (*R²* = 0.99; Fig. 8c).

**Figure 8:**
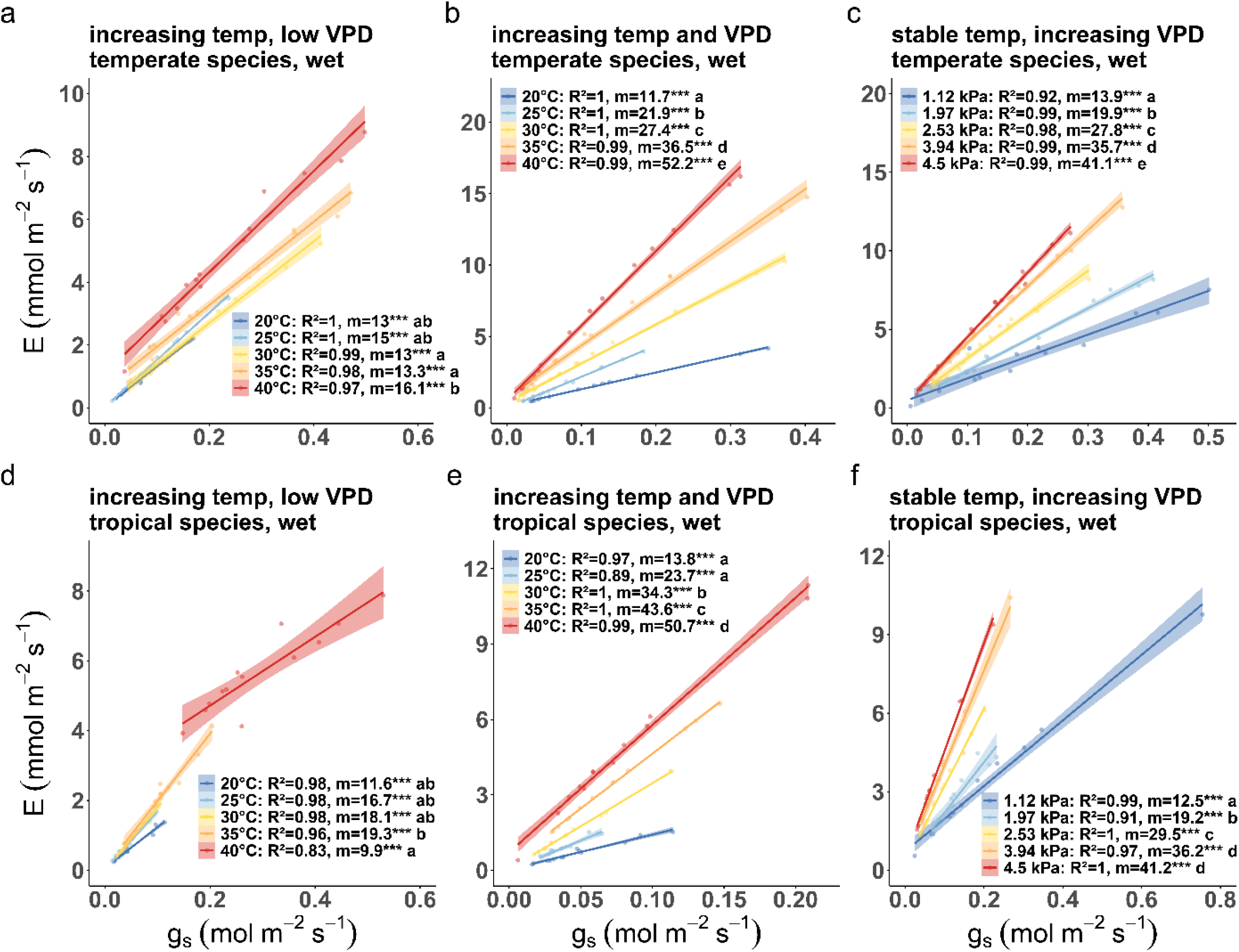
Changes in the relationship between transpiration (E) and stomatal conductance (g_s_) when temperature (T_air_) increases while vapour pressure deficit (VPD) remains low (a, d), T_air_ and VPD are simultaneously increasing (b, e), and VPD increases at a stable T_air_ of 35°C (c, f) in temperate (a, b, c) and tropical (d, e, f) tree species of well-watered plants. Points show individual measurements; colored lines are linear fits plotted only when slopes are significant (p < 0.05) from global interaction models (A_net_ ∼ g_s_ × Temp or A_net_ ∼ g_s_ × VPD), with shaded 95% confidence intervals. Legends report per-level R² and slope m (A_net_/g_s_) with significance asterisks (* p < 0.05, ** p < 0.01, *** p < 0.001), and compact letter displays summarize pairwise slope differences.

In tropical species, E*–*g_s_ correlations were equally strong but displayed slightly lower *R²* values at the highest T_air_ under low VPD (Fig. 8d–f). Under low VPD, slopes varied between 11.6*** (ab) at 20°C and 9.9*** (a) at 40°C (*R²* = 0.83–0.98; Fig. 8d). When T_air_ and VPD increased together, *m* rose steadily from 13.8*** (a) at 20°C (*R²* = 0.97) to 50.7*** (d) at 40°C (*R²* = 0.99; Fig. 8e). At constant T_air_, increasing VPD again resulted in steeper slopes, with *m* = 12.5*** (a) at 1.12 kPa (*R²* = 0.99) and *m* = 41.2*** (d) at 4.5 kPa (*R²* = 1; Fig. 8f).

Under drought, the E*–*g_s_ relationship remained nearly linear but shifted toward steeper slopes, indicating higher transpirational output per unit stomatal conductance (Fig. S25). In temperate species, *R²* values stayed ≥ 0.89 across all T_air_. Slopes increased from 13.7*** (a) at 20°C to 25.6*** (a) at 40°C (Fig. S25a). When T_air_ and VPD rose simultaneously, *m* increased strongly from 13.1*** (a) at 20°C to 63*** (e) at 40°C (*R²* = 0.99–1; Fig. S25b). At constant T_air_, VPD caused a similar monotonic steepening: *m* = 16.4*** (a) at 1.12 kPa to 47.5*** (e) at 4.5 kPa (*R²* = 0.96–1; Fig. S25c).

Drought-exposed tropical species also exhibited near-perfect E*–*g_s_ coupling (Fig. S25d–f). At low VPD, slopes increased modestly with T_air_, from 18.1*** (a) at 25°C (*R²* = 0.98) to 16.2*** (a) at 40°C (*R²* = 0.90; Fig. S25d). When T_air_ and VPD rose together, slopes steepened sharply, from 13.9*** (a) at 20°C to 65.6*** (d) at 40°C (*R²* = 0.94–1; Fig. S25e). Under constant 35°C, the relationship again strengthened with VPD, with *m* = 16.2*** (a) at 1.12 kPa (*R²* = 0.98) and 44.8*** (e) at 4.5 kPa (*R²* = 1; Fig. S25f).

Overall, E*–*g_s_ coupling remained almost perfectly linear across treatments, confirming the dominant control of stomatal conductance on transpirational water loss. The progressively steeper slopes with an increasing VPD, both when T_air_ increased or remained stable at 35°C, demonstrate the enhanced water flux per unit stomatal conductance. In conclusion, when T_air_ increases under a constant low VPD, E can only be increased by increasing g_s_ (Figs. 8a, 8d, S25a, S25d), while under simultaneous increasing VPD, g_s_ can remain low (Figs. 8b, 8e, S25b, S25e), or decrease when VPD increased at a stable T_air_ (Figs. 8c, 8f, S25c, S25f) to maintain E.

### T_air_ and VPD effects on leaf temperature

The temperature difference between leaves and air (T_offset_; T_leaf_ – T_air_) decreased with rising T_air_ and VPD in both biogeographic groups, indicating enhanced evaporative cooling under high thermal load (Figs. S26–S28).

Under well-watered conditions and low VPD, temperate species exhibited no BP (Davies *p* = 0.285) but showed a steady decline in T_offset_ from +0.13 ± 0.06°C at 20°C to –0.16 ± 0.09°C at 40°C (*R²* = 0.11, *p* = 0.0042, slope = –0.014; Fig. S27a). Tropical species displayed a BP at 31.29°C (Davies *p* = 0.012): below this threshold, T_offset_ remained stable (+0.55 ± 0.19°C at 20°C to +0.79 ± 0.19°C at 30°C), whereas above it, T_offset_ declined (*R²* = 0.28, *p* = 0.0074, slope = –0.108), reaching –0.11 ± 0.11°C at 40°C (Fig. S27b). When T_air_ and VPD increased simultaneously, both groups exhibited continuous decreases (Davies *p* = 0.543, *R²* = 0.21, *p* = 6.2 × 10⁻⁵, slope = –0.056, and Davies *p* = 0.0815, in temperate and tropical species, respectively. Fig. S27c, d). At constant 35°C, T_offset_ declined with increasing VPD in temperate species (*R²* = 0.29, *p* = 6.8 × 10⁻⁷, slope = –0.224; Fig. S27e) from +0.28 ± 0.11°C at 1.12 kPa to –0.50 ± 0.15°C at 4.5 kPa, whereas tropical species exhibited a BP at 3.22 kPa (Davies *p* = 0.0354): below this threshold, T_offset_ increased slightly (*R²* = 0.14, *p* = 0.033, slope = 0.282), while above it, no further change was detected (*R²* = 0.06, *p* = 0.23, slope = –0.550; Fig. S27f).

Under drought, T_offset_ responses were largely attenuated (Fig. S28). No BPs were detected for either species group when T_air_ increased (Davies *p* > 0.05; Fig. S28a–d). At constant T_air_, temperate species showed a gradual decrease from +0.60 ± 0.10°C at 1.12 kPa to +0.33 ± 0.08°C at 4.5 kPa (Davies *p* = 0.532; Fig. S28e). In tropical species, a BP occurred at 3.41 kPa (Davies *p* = 0.00097): below this point, T_offset_ increased (*R²* = 0.27, *p* = 0.0015, slope = 0.401), reaching +0.94 ± 0.16°C at 2.53 kPa, whereas above it, T_offset_ decreased (*R²* = 0.19, *p* = 0.028, slope = –0.719), falling to +0.43 ± 0.09°C at 4.5 kPa (Fig. S28f).

### T_air_ and VPD effects on leaf temperature in the dark

While T_leaf_ in the dark was consistently below T_air_, the trends in leaf–to-air temperature offsets in darkness (T_offset dark_, Figs. S29–S31) were comparable to those observed in the light.

Under wet conditions, T_offset dark_ of temperate species decreased steadily from –0.67 ± 0.07°C at 20°C to –1.27 ± 0.16°C at 40°C (Fig. S30a). Tropical species displayed a BP at 33.2°C (Davies *p* = 0.0139): below this point, no response was detected (*R²* = 0.00, *p* = 0.71, slope = 0.003), whereas above it, T_offset dark_ decreased significantly (*R²* = 0.22, *p* = 0.022, slope = –0.053), from –0.65 ± 0.04°C at 30°C to –0.99 ± 0.12°C at 40°C (Fig. S30b). When T_air_ and VPD rose together, both species groups exhibited continuous cooling (*R²* = 0.13, *p* = 0.0021, slope = –0.038, and Davies *p* = 0.00217, *R²* = 0.58, *p* = 5.9 × 10⁻⁶, slope = –0.148 in temperate and tropical species, respectively. Fig. S30c, d). At constant 35°C, T_offset dark_ showed small VPD effects, declining from –0.86 ± 0.11°C at 1.12 kPa to –1.16 ± 0.21°C at 4.5 kPa in temperate species (*R²* = 0.06, *p* = 0.032, slope = –0.128; Fig. S30e), while tropical species remained stable (Davies *p* > 0.05; Fig. S30f).

Under drought, T_offset dark_ remained constant across all treatments (Fig. S31a, b, c, d, f), except in temperate species at constant 35°C with rising VPD, with a slight significant increase in cooling (–0.60 ± 0.08°C at 1.12 kPa to –0.82 ± 0.11°C at 4.5 kPa; *R²* = 0.08, *p* = 0.015, slope = –0.085; Fig. S31e).

Overall, under a sufficient water supply, T_offset_ and T_offset dark_ decreased with increasing T_air_, but much more efficiently when VPD increased simultaneously (Figs. S26–S31). Despite this general trend, clear biogeographic contrasts emerged. Temperate species exhibited a stronger cooling response and larger absolute decreases in T_offset_ with increasing T_air_ and VPD than tropical species when light-exposed (Fig. S27). In contrast, T_leaf_ of tropical species was largely higher relative to T_air_, with even delayed declines in T_offset_, indicating stronger radiative energy uptake. Under drought, evaporative cooling weakened in both groups, and no effective cooling of T_leaf_ below T_air_ remained.

### The relationship between leaf temperature offsets and stomatal conductance, and transpiration, with and without light

Both with and without light, T_offset_ became increasingly smaller (cooler leaves) with rising g_s_ and E (Figs. S32–S39), confirming overall that evaporative fluxes exerted a strong physical influence on T_leaf_. However, the strength and consistency of this coupling were much stronger in temperate than in tropical species and weakened under drought.

Under well-watered conditions, temperate species exhibited tight and mostly linear T_offset_–g_s_ relationships, particularly when T_air_ and VPD rose simultaneously. Slopes became increasingly negative with rising T_air_, reaching *m* = –8.11*** at 35°C and *m* = –7.72*** at 40°C (*R*² = 0.78 and 0.71; Fig. S32b), with cooling apparent once g_s_ exceeded 0.05 mol m⁻² s⁻¹. At constant 35°C, rising VPD further strengthened this relation (*R*² = 0.62, *m* = –3.36***; Fig. S32c). In tropical species T_offset_–g_s_ correlations were much weaker and less consistent, with significant cooling only at the highest T_air_ or VPD (e.g., *R*² = 0.73, *m* = –13.07**, Fig. S32e; *R*² = 0.57, *m* = –8.45***, Fig. S32f), and cooling occurring at higher g_s_ thresholds (> 0.18 mol m⁻² s⁻¹).

Under drought, T_offset_–g_s_ coupling weakened substantially (Fig. S33). Significant relations persisted only in a few cases, such as temperate leaves at 35°C (*R*² = 0.34, *m* = –12.85**, Fig. S33a) and tropical leaves under combined heat and VPD (*R*² = 0.46, *m* = –5.96*, Fig. S33d). Overall slopes across all T_air_ were non-significant for most treatments, suggesting that the link between stomatal opening and leaf cooling was strongly limited once water supply was limited.

In darkness, a consistent T_offset_*–*g_s_ relationship within the treatments was restricted to the temperate species under high T_air_ under well-watered conditions (Fig. S34). When T_air_ increased, temperate species displayed overall a consistent linear decrease in T_offset dark_ with increasing g_s dark_ (across all species: *R*² = 0.47, *m* = –2.75, *p* = 1 × 10⁻¹⁰ and *R^2^* = 0.44, *m* = -6.64, p = 2.9 × 10⁻¹⁰, respectively. Fig. S34a, b). However, this relationship was only significant at 30°C and above. In contrast, while the overall relationship across all T_air_ or VPD remained in tropical species (overall *R*² = 0.43, *m* = –2.45, *p* = 7.9 × 10⁻^9^; and *R*² = 0.24, *m* = –7.2, *p* = 4.1 × 10⁻^5^. Fig. S34d, e), it was inconsistent and often absent within a given T_air_, especially when VPD increased (Fig. S34e). At constant T_air_, increasing VPD accentuated cooling responses in temperate species (e.g., *R*² = 0.84, *m* = –11.08*** at 4.5 kPa, Fig. S34c), but again, this relationship was much less consistent among the tropical species (Fig. S34f). Under drought (Fig. S35), however, these relations became sporadic and inconsistent across T_air_ steps, with significant trends confined to tropical species when T_air_ increased at a low VPD (*R*² = 0.52, *m* = –2.97, *p* = 1.5 × 10⁻¹⁰; Fig. S35d).

The decline of T_offset_ with increasing E was mainly visible across all T_air_ and VPD (Figs. S36–S39). In well-watered plants, the T_offset_–E relationship was particularly strong in temperate species when T_air_ and VPD increased together (*R*² = 0.76, *m* = –0.18, *p* = 2.9 × 10⁻²³; Fig. S36b), indicating that an increase in E yielded in actual leaf cooling (T_offset_ < 0 when E > 1.9 mmol m⁻² s⁻¹). Tropical species showed weaker associations (*R*² = 0.45, *m* = –0.22, *p* = 7.9 × 10⁻¹⁰; Fig. S36e), requiring nearly doubled transpiration rates (> 3.7 mmol m⁻² s⁻¹) to achieve comparable cooling.

In darkness, and therefore in the absence of interferences by light-energy inputs of the leaf energy balance, the relationships between T_offset dark_ and E_dark_ were again only more consistently significant when all measurements were included across all T_air_ and VPD treatments (*R*² = 0.33–0.48, *m* ≈ – 0.14 to –0.20, *p* ≤ 1.26 10⁻6; Figs. S38). A significant relationship between T_offset dark_ and E_dark_ within the different T_air_ or VPD steps was again inconsistent, and largely restricted to temperate species at high T_air_. The slopes flattened or reversed under drought (Figs. S39), further revealing a strong reduction in the efficiency of evaporative cooling once hydraulic limitation set in.

Taken together, T_offset_ declined only overall across all T_air_ by increasing g_s_ and E, but not consistently within the separate T_air_ steps, where E effectively contributes to leaf cooling mostly at high T_air_ ≥ 30°C in temperate but not in tropical species when well-watered (Figs. S32, S34, S36, S38). Interestingly, leaf T_leaf_ of tropical species was consistently higher than that of temperate species. This difference cannot be attributed to reduced evaporative cooling efficiency, as the overall relationship (slopes) between T_offset_ and E (Fig. S36, S38) was similar. Instead, tropical leaves appear to operate at a higher thermal equilibrium when exposed to light, but not during darkness (Figs. S26, S29), implicating greater radiative heat absorption (e.g., higher short-wave absorptance or differences in wax/roughness affecting absorptance/emissivity and the near-leaf boundary layer) as the primary driver.

## Discussion

### The decoupling of photosynthesis from stomatal conductance at high T_air_

Across treatments, we observed a T_air_-driven divergence and a VPD-driven convergence between photosynthetic assimilation (Aₙₑₜ) and stomatal conductance (gₛ), arising from the joint influence of T_air_ and atmospheric demand, and further modulated by soil moisture availability (Figs. 1, 2, 6, S1, S2, S3, S4, S23).

Within the tested T_air_ range (20–40°C), a T_air_-driven stomatal decoupling, here defined as a constantly decrease or loss of the relationship between A_net_ and g_s_ with rising T_air_, could only be observed in temperate species when VPD was kept low (Fig. 6). As T_air_ increased at low VPD, A_net_ rose only up to approximately 35°C and then levelled or declined (Fig. 1a; S1a), whereas g_s_ continued to increase (Fig. 2a; S3a), resulting in a progressive flattening of the A_net_-g_s_ relationship (i.e., the slope (m) from 56.3 at 20°C to 20.5 at 40°C (R² = 0.85–0.79; Fig. 6a). Since the leaf-to-air vapor gradient remained too small to increase E efficiently (Fig. 8a), plants had to actively raise g_s_ to increase transpirational flux, especially at high T_air_ above 30°C (E, Figs. 3a, S11), similar to what has been observed by Diao et al., 2024a. Furthermore, the strong and highly linear relationships between E and g_s_ (Figs. 8, S25) confirm that stomatal regulation remained fully operational even at extreme conditions. Thus, it is implausible that the observed decoupling results from a breakdown of stomatal control but rather from an active upregulation of g_s_ and therefore E, despite thermal constraints on photosynthesis. This agrees with the findings of Wang et al. 2024, where the temperature-driven breakdown of the minimum leaf conductance occurred at T_air_ > 40°C. In contrast, when T_air_ and VPD increased together, we could not observe a decoupling between A_net_ and g_s_, since the relationship between A_net_ and g_s_ remained largely unchanged (Fig. 6b; S23b), further demonstrating an effective stomatal control within the tested temperature range. The E increased, even if g_s_ changed little or declined (Fig. 2b; S3d; S4d; S11c, d), since the rising VPD directly amplifies transpiration, as further reflected in the steepening E-g_s_ relationships with increasing VPD (Fig. 8b). A similar response was observed at constant 35°C with rising VPD, similar to Diao et al., 2024b, where g_s_ remained stable or decreased (Fig. 2c; S3e; S4e), while E increased (S11e), again showing that rising VPD maintained a constant E while g_s_ decreased (Fig. 8c).

Interestingly, in tropical species, the coupling between A_net_ and g_s_ (and hence between A_net_ and E) was weak or even absent at low VPD across much of the T_air_ range (Fig. 6d; Figs. 7d; S23d; S24d), but re-emerged when VPD increased, either alongside T_air_ or at a constant high Tair (Figs. 6e, f; Fig. 7e, f; S23e, f; S24e, f). Drought further increased the coupling between A_net_ and g_s_, reflecting overall what has been seen in trees in the Amazon rain forest (Janssen et al., 2020). The largely absence of coordination between stomatal conductance and photosynthesis at low VPD might derive from their adaptation to monsoonal climates, where low VPD and abundant water availability during the rainy season do not require a conservative, water-saving stomatal regulation, whereas VPD increases along with decreased water availability during the dry season, therefore fostering a more water-conservative stomatal control. A stronger sensitivity of g_s_ to VPD than drought (Figs. 2d, e, S4b, d) has been observed in a tropical rainforest in Panama during an El Niño event (Wu et al., 2020).

Earlier studies reported an Aₙₑₜ-gₛ decoupling when both Tₐᵢᵣ and VPD increased (Marchin et al., 2023; Gauthey et al., 2024), but those cases typically involved leaf T_air_ exceeding 40–45°C, where photosynthetic biochemistry collapses (Medlyn et al., 2002; Didion-Gency et al., 2025) while stomata remain partially open or leaf physical properties are no longer able to prevent a runaway water loss (Riederer and Schreiber, 2001; Wang et al., 2024). In contrast, in our tested range (≤ 40°C), photosynthesis persisted, and gₛ adjustments remained functional, indicating that the observed decoupling arises from a true physiological mechanism for fluxes regulations rather than from thermal damage of the stomatal apparatus.

The observed rise in E under high VPD, despite partial stomatal closure, underscores the dominant role of evaporative demand in driving water loss. Because E ≈ gₛ × ΔVPD (where ΔVPD is the leaf–air vapor gradient), E may increase either through higher gₛ or through a larger ΔVPD. When VPD remains low, gₛ must rise to elevate E; when VPD increases, ΔVPD dominates, allowing gₛ to remain constant or decline while E still rises (Figs. 1–2; S11–S12). Thus, the Aₙₑₜ-gₛ decoupling at constant low VPD is the hydraulic counterpart of an attempt to sustain transpirational flux under thermal stress. This mechanistic interpretation is consistent with the contrasting behaviors of intrinsic (WUEᵢ = Aₙₑₜ/gₛ) and instantaneous (WUE = Aₙₑₜ/E) water-use efficiencies. At constant VPD, increasing Tₐᵢᵣ caused WUEᵢ and WUE to decline sharply (Figs. S19–S22), reflecting the steep rise in gₛ (and E) relative to the modest gain or decline in Aₙₑₜ. In contrast, when VPD increased with T_air_, WUE continued to decline even though gₛ remained stable, because E rose through the enhanced VPD gradient. Together, these responses show that the Aₙₑₜ–gₛ decoupling is a consequence of maintaining transpirational flux under high T_air_.

Lastly, the near lack of changes in the ratio between gₛ and _gs dark_, as well as E and E_dark_ (Figs. S8- S10, S16-S18), indicates a common control. Interestingly, their decrease in temperate species under a constant low VPD and rising T_air_ was driven by the much stronger increase in g_s dark_ than the observed increase in g_s_, leading them to be very similar at 40°C (Fig. S9a). This finding challenges the assumption that stomatal regulation is primarily light-driven, at least at high T_air_ ≥ 30°C. Hence, the term “residual conductance” may underestimate the functional role of gₛ _dark_ in maintaining continuous water flow, and therefore, other services such as nutrient and O_2_/CO_2_ transport in heterotrophic tissues such as roots and stems (Levy et al., 1999; Sorz and Hietz, 2006; Jensen et al., 2016).

### Leaf thermal regulation

From a thermal perspective, T_air_-driven decoupling enhances transpiration and should therefore contribute to evaporative cooling (Gauthey et al., 2024; Bachofen et al., 2025), and we found a consistent cooling of T_leaf_ below T_air_ only in temperate species (Figs. S26-S28). However, the relationships between T_offset_ and g_s_ and T_offset_ and E are only consistently significant in temperate species in high temperatures (Figs. S26–S39). Furthermore, likely because the thermal conductivity of air decreases with increasing humidity (Pernau et al., 2024), the relationship between E and leaf cooling was very low under the conditions we actually observed stomatal decoupling (Fig. S36). Across all T_air_ and VPD steps, temperate and tropical species achieved a similar overall cooling efficiency per unit E (Figs. S36-S39). However, tropical species had leaves that were consistently warmer than leaves of temperate species and were rarely below T_air_, something that has been observed before (Crous et al., 2023; Middleby et al., 2025a, 2025b). Since the lower cooling ability with rising E of tropical species remains in the dark (Figs. S38), this points to a combination of radiative and structural differences (e.g., surface absorptance, boundary-layer conductance) as primary determinants of T_leaf_–T_air_ offsets (Kullberg and Feeley, 2024; Middleby et al., 2025a).

Overall, leaf transpirational cooling, at least in the investigated species, appears rather as a physical consequence of sustained transpiration rather than a targeted outcome of stomatal control and therefore, the observed decoupling.

## Conclusions and implications

In conclusion, (1) we could confirm that stomatal regulation at high T_air_, and therefore stomatal decoupling, will be strongly impacted by VPD and soil moisture availability, with reductions in g_s_ at high VPD and in dry soil, while rising T_air_ at stable low VPD and moist soil conditions leads to stomatal decoupling and increased E. Interestingly, we found that (2) tree species from temperate regions had a stronger stomatal regulation across T_air_ ranges, with a decoupling at high T_air_, low VPD, and moist conditions. In contrast, tropical species, which evolve at higher T_air_ and VPD environments, only barely co-regulated A_net_ and g_s_ at low VPD. Furthermore, (3) we found that the increase of E by stomatal decoupling only barely contributed to T_leaf_ regulation, while a VPD increase was much more efficient due to a better heat flux.

Taken together, our results indicate that stomatal decoupling represents a biophysical strategy to sustain transpirational water flux under high T_air_, not a failure of regulation nor an active optimization for cooling. When Tₐᵢᵣ increases under constant VPD, gₛ must rise to elevate E (Fig. 8a), leading to a decrease in the relationship between A_net_ and g_s_, and therefore a strong decline in WUE_i_ (Fig. S19a) as well as WUE (Fig. S21a). When VPD increases, ΔVPD drives E (Fig. 8b), preserving Aₙₑₜ–gₛ coupling and therefore a stable WUE_i_ (Fig. S19c) but reducing WUE (Fig. S21c) through atmospheric control.

These findings could be important at the parameter level to model g_s_, as it points out that the *g_0_* parameter, for which g_s dark_ can be taken as a proxy, representing the “residual” stomatal opening independent of photosynthetic activity (Medlyn et al., 2011). In temperate species, *g₀* (as approximated by g_s dark_) slightly increased with Tair under well-watered, low-VPD conditions, and we found a clear breakpoint with a much steeper increase around 35°C (Fig. S6a). In tropical species, a similar change in the relationship occurred at slightly lower T_air_ (∼33°C; Fig. S6b). When T_air_ and VPD increased simultaneously, g_s dark_ remained largely unchanged in both groups (Figs. S6c, d), and declined sharply when VPD increased at a constant 35°C (Figs. S6e, f). Under drought, however, _gs dark_ increased only at a low rate with rising T_air_ at a low VPD (Figs. S7a, b), and exhibited breakpoint-type accelerated decline with rising VPD (Figs. S7e, f), demonstrating a stronger regulation of the stomatal conductance during darkness in response to reduced water availability. Together, these trends suggest that *g₀* is not a fixed trait but a dynamic quantity that increases with T_air_ at a low evaporative demand independent of water limitations, yet is stable or decreases when VPD rises. Together, these findings demonstrate that g₀ should not be seen as a fixed residual conductance, especially not at high T_air_, but a dynamic physiological parameter actively modulated by T_air_, VPD, and water availability. This suggests that stomata at high T_air_ remain more open than expected, even in the dark, to maintain a continuous water flow through the plant, supporting hydraulic and possibly other plant metabolic demands. Furthermore, similar to Middleby et al., 2024, we found further support that the *g_1_* parameter in the Medlyn model (Medlyn et al., 2011), which is inversely related to WUE_i_, changes with T_air_, VPD, and soil moisture availability (Figs. S19, S20), including distinct T_air_ breaking points above which its trend reverses, both in temperate and tropical species.

Other studies will need to investigate the behavior of stomatal regulation at more extreme T_air_ above 40°C, where the loss of physiological control, for instance through increased cuticular permeability (Riederer and Schreiber, 2001; Wang et al., 2024), and biochemical dysfunctionality (Gauthey et al., 2024; Didion-Gency et al., 2025), will likely become more important.

## SNAcknowledgments

PS, MD-G, TJ, CB, GB, and CG were supported by the Swiss National Science Foundation SNSF (310030_204697 and CRSK-3_220989) and the Sandoz Family Foundation. We thank Georges Grun for his support with the climate chamber control.

## Author contribution

PS and CG planned and designed the research. PS, MD-G, TJ, and CG conducted the laboratory work. GH and AK maintained the climate chamber facility. PS analyzed the data and led the manuscript’s writing, and PS, CG, MD-G, TJ, GH, AK, CB, and GB contributed to the final version.

## Data availability

Upon the manuscript’s acceptance, the data supporting the findings will be made openly available in a public repository (Envidat).

## Supporting information

Overview Main and Supplementary Figures

## Supplementary tables

**Table S1:** Characteristics of the studied tree species.

| Latin name | Family | Natural range | Climate<br>(Köppen) | Leaf habit |
| --- | --- | --- | --- | --- |
| *Alnus cordata*<br>(Loisel.) Duby | Betulaceae | Italy | Temperate<br>(Cfa – Cfb) | Deciduous |
| *Acer platanoides* L. | Sapindaceae | Europe | Temperate<br>(Cfa) | Deciduous |
| *Phillyrea angustifolia*<br>L. | Oleaceae | Southern Europe to<br>Northern Africa | Temperate<br>(Csa) | Evergreen |
| *Terminalia*<br>*microcarpa* Decne. | Combretaceae | Southeast Asia to<br>North Australia | Tropical<br>(Am-Af) | Semi-deciduous |
| *Trema tomentosa*<br>(Roxb.) H. Hara | Cannabaceae | South and<br>Southeast Asia to<br>Australia | Tropical<br>(Am-Af) | Evergreen |
| *Syzygium jambos* L.<br>(Alston) | Myrtaceae | Southeast Asia | Tropical<br>(Am-Af) | Evergreen |

**Table S2:** Average soil moisture (SM, volume-%), the corresponding standard errors for the overall treatments, as well as for the single days of the experiment. The final trend analysis (trend LM) indicates if there was a significant change in SM during the experimental period.

|  | increasing temp, low VPD |  |  |  | increasing temp and VPD |  |  |  | constant temp,increasing VPD |  |  |  |
| --- | --- | --- | --- | --- | --- | --- | --- | --- | --- | --- | --- | --- |
|  | wet |  | dry |  | wet |  | dry |  | wet |  | dry |  |
|  | SM % | SE | SM % | SE | SM % | SE | SM % | SE | SM % | SE | SM % | SE |
| **overall** | 32.6 | 0.3 | 9.1 | 0.3 | 32.4 | 0.4 | 9.15 | 0.3 | 31.2 | 0.4 | 7.4 | 0.3 |
| **2** | 33.8 | 0.9 | 12.2 | 0.7 | 34.4 | 0.9 | 12.7 | 0.9 | 35.2 | 0.6 | 11.7 | 0.8 |
| **4** | 33.3 | 0.7 | 10.4 | 0.7 | 31.3 | 1.1 | 10.3 | 0.8 | 30.2 | 0.8 | 8.1 | 0.7 |
| **6** | 32.6 | 0.8 | 9.3 | 0.7 | 32.3 | 0.7 | 9.4 | 0.8 | 29.0 | 1.0 | 6.7 | 0.5 |
| **8** | 32.8 | 0.8 | 8.2 | 0.6 | 31.5 | 1.0 | 8.3 | 0.7 | 29.0 | 1.2 | 6.2 | 0.4 |
| **10** | 29.8 | 0.8 | 7.3 | 0.5 | 32.1 | 0.8 | 7.7 | 0.6 | 32.0 | 0.9 | 6.1 | 0.3 |
| **12** | 33.4 | 0.7 | 7.1 | 0.5 | 32.6 | 1.0 | 6.6 | 0.6 | 31.6 | 0.9 | 5.6 | 0.2 |
|  | **R<sup>2</sup>** | **p** | **R<sup>2</sup>** | **p** | **R<sup>2</sup>** | **p** | **R<sup>2</sup>** | **p** | **R<sup>2</sup>** | **p** | **R<sup>2</sup>** | **p** |
| **trend (LM)** | 0.02 | 0.069 | 0.224 | 8.90E-11 | 0.006 | 0.338 | 0.194 | 2.50E-09 | 0.014 | 0.133 | 0.294 | 3.20E-14 |

**Table S3:** Mean measured Tair and VPD with the corresponding standard errors in the climate chambers and the measurement chamber of the Li-6800.

| increasing temperature, low VPD |  |  |  |  |  |  |  | increasing temperature and VPD |  |  |  |  |  |  |  | stable temperature, increasing VPD |  |  |  |  |  |  |  |
| --- | --- | --- | --- | --- | --- | --- | --- | --- | --- | --- | --- | --- | --- | --- | --- | --- | --- | --- | --- | --- | --- | --- | --- |
| Chamber |  | Li-6800 |  | Chamber |  | Li-6800 |  | Chamber |  | Li-6800 |  | Chamber |  | Li-6800 |  | Chamber |  | Li-6800 |  | Chamber |  | Li-6800 |  |
| Temp | Temp | Temp | Temp | VPD | VPD | VPD | VPD | Temp | Temp | Temp | Temp | VPD | VPD | VPD | VPD | Temp | Temp | Temp | Temp | VPD | VPD | VPD | VPD |
| (°C) | +/- SE | (°C) | +/- SE | (kPa) | +/- SE | (kPa) | +/- SE | (°C) | +/- SE | (°C) | +/- SE | (kPa) | +/- SE | (kPa) | +/- SE | (°C) | +/- SE | (°C) | +/- SE | (kPa) | +/- SE | (kPa) | +/- SE |
| 19.8 | 0.04 | 20.1 | 0.065 | 1.2 | 0.003 | 1.18 | 0.005 | 19.5 | 0.070 | 20.3 | 0.111 | 1.1 | 0.010 | 1.2 | 0.011 | 34.8 | 0.045 | 35.0 | 0.006 | 1.4 | 0.020 | 1.6 | 0.036 |
| 25.0 | 0.04 | 25.3 | 0.070 | 1.4 | 0.005 | 1.34 | 0.017 | 24.9 | 0.029 | 25.3 | 0.105 | 2.0 | 0.005 | 2.1 | 0.017 | 35.0 | 0.023 | 35.0 | 0.003 | 2.2 | 0.007 | 2.0 | 0.035 |
| 30.0 | 0.03 | 30.0 | 0.006 | 1.4 | 0.006 | 1.37 | 0.015 | 30.0 | 0.013 | 30.0 | 0.004 | 3.1 | 0.004 | 3.1 | 0.005 | 35.0 | 0.015 | 35.0 | 0.009 | 3.0 | 0.004 | 3.0 | 0.020 |
| 35.1 | 0.03 | 35.0 | 0.005 | 1.5 | 0.008 | 1.68 | 0.019 | 35.0 | 0.015 | 35.0 | 0.006 | 4.5 | 0.006 | 4.5 | 0.005 | 35.0 | 0.014 | 35.0 | 0.005 | 4.0 | 0.003 | 4.0 | 0.021 |
| 38.8 | 0.20 | 40.0 | 0.006 | 1.9 | 0.010 | 2.13 | 0.043 | 38.8 | 0.197 | 40.0 | 0.007 | 5.6 | 0.057 | 6.0 | 0.005 | 34.1 | 0.131 | 35.0 | 0.003 | 4.3 | 0.039 | 4.6 | 0.010 |

## Notes

### Competing Interest Statement

The authors have declared no competing interest.

