## Supplementary material for "Decoupling of stomatal conductance from net assimilation at high temperature as a mechanism to increase transpiration": Overview Main and Supplementary Figures

### Main Figures

**1: $A_{net}$ - Overview temperature and VPD response**

**2: $g_s$ - Overview temperature and VPD response**

**3: E - Overview temperature and VPD response**

**4: $WUE_I$ - Overview temperature and VPD response**

**5: WUE - Overview temperature and VPD response**

**6: $A_{net}$ vs. $g_s$, wet**

**7: $A_{net}$ vs. E, wet**

**7: E vs. $g_s$, wet**

### SI Figures

**S1: $A_{net}$ - Temperature and VPD response, including breaking point analysis, wet**

**S2: $A_{net}$ - Temperature and VPD response, including breaking point analysis, dry**

**S3: $g_s$ - Temperature and VPD response, including breaking point analysis, wet**

**S4: $g_s$ - Temperature and VPD response, including breaking point analysis, dry**

**S5: $g_{s\ dark}$ - Overview temperature and VPD response**

**S6: $g_{s\ dark}$ - Temperature and VPD response, including breaking point analysis, wet**

**S7: $g_{s\ dark}$ - Temperature and VPD response, including breaking point analysis, dry**

**S8: $g_s$-ratio ($g_s$ divided by $g_{s\ dark}$) - Overview temperature and VPD response**

**S9: $g_s$-ratio - Temperature and VPD response, including breaking point analysis, wet**

**S10: $g_s$-ratio - Temperature and VPD response, including breaking point analysis, dry**

**S11: E - Temperature and VPD response, including breaking point analysis, wet**

**S12: E - Temperature and VPD response, including breaking point analysis, dry**

**S13: $E_{dark}$ - Overview temperature and VPD response**

**S14: $E_{dark}$ - Temperature and VPD response, including breaking point analysis, wet**

**S15: $E_{dark}$ - Temperature and VPD response, including breaking point analysis, dry**

**S16: E-ratio (E divided by $E_{dark}$) - Overview temperature and VPD response**

**S17: E-ratio - Temperature and VPD response, including breaking point analysis, wet**

**S18: E-ratio - Temperature and VPD response, including breaking point analysis, dry**

**S19: $WUE_I$ - Temperature and VPD response, including breaking point analysis, wet**

**S20: $WUE_I$ - Temperature and VPD response, including breaking point analysis, dry**

**S21: WUE - Temperature and VPD response, including breaking point analysis, wet**

**S22: WUE - Temperature and VPD response, including breaking point analysis, dry**

**S23: $A_{net}$ vs. $g_s$, dry (aka Fig. 6 in dry)**

**S24: $A_{net}$ vs. E, dry (aka Fig. 7 in dry)**

**S25: E vs. $g_s$, dry (aka Fig. 8 in dry)**

**S26: $T_{offset}$ - Overview temperature and VPD response**

**S27: $T_{offset}$ - Temperature and VPD response, including breaking point analysis, wet**

**S28: $T_{offset}$ - Temperature and VPD response, including breaking point analysis, dry**

**S29: $T_{offset\ dark}$ - Overview temperature and VPD response**

**S30: $T_{offset\ dark}$ - Temperature and VPD response, including breaking point analysis, wet**

**S31: $T_{offset\ dark}$ - Temperature and VPD response, including breaking point analysis, dry**

**S32: $T_{offset}$ vs $g_s$, wet**

**S33: $T_{offset}$ vs $g_s$, dry**

**S34: $T_{offset\ dark}$ vs $g_{s\ dark}$, wet**

**S35: $T_{offset\ dark}$ vs $g_{s\ dark}$, dry**

**S36: $T_{offset}$ vs E, wet**

**S37: $T_{offset}$ vs E, dry**

**S38: $T_{offset\ dark}$ vs $E_{dark}$, wet**

**S39: $T_{offset\ dark}$ vs $E_{dark}$, dry**
